# A dual mechanism of APC/C inhibition by MAP kinases

**DOI:** 10.1101/2022.04.01.485959

**Authors:** Li Sun, Shuang Bai, Jia-li Chen, Da-jie Deng, Zhou-qing Luo, Yamei Wang, Quan-wen Jin

## Abstract

Mitotic anaphase onset is a key cellular process that is tightly regulated by multiple kinases. The involvement of mitogen-activated protein kinases (MAPKs) in this process has been established long ago in *Xenopus* egg extracts. However, despite its importance, it is still unclear which MAPK(s) is actually involved, this impedes the further understanding of the regulatory cascade. In this study, we first demonstrated that the involvement of MAPKs in mitotic anaphase onset regulation is evolutionarily conserved in the fission yeast (*Schizosaccharomyces pombe*). Then, we found that two of the three fission yeast MAPK signaling pathways act in concert to restrain anaphase-promoting complex/cyclosome (APC/C) activity upon activation of the spindle assembly checkpoint (SAC). The first pathway involves the phosphorylation of Mad2, a component of the core mitotic check complex (MCC), by MAPK Sty1, which enhances the tight binding of MCC to APC/C. The second pathway involves MAPK Pmk1 phosphorylation of Slp1^Cdc20^, the fission yeast homologue of Cdc20 and the co-activator of APC/C, which promotes the degradation of Slp1^Cdc20^. Both phosphorylation events are required to sustain mitotic arrest in response to spindle defects. These results clarified a detailed regulation cascade of the ubiquitous MAPK signaling in spindle checkpoint activation, APC/C inhibition and anaphase entry, which is vital for accurate chromosome segregation and cell viability.

## INTRODUCTION

The evolutionarily conserved mitogen-activated protein kinase (MAPK) signaling pathways regulate multiple cellular functions in eukaryotic organisms in response to a wide variety of environmental cues (Plotnikov et al., 2011). However, different MAPK pathways have been evolved in an organism to integrate wide range of signals and fulfill different regulations on various effectors (Cansado et al., 2021; Ronkina and Gaestel, 2022). This makes specifying the MAPKs and substrates involved in a specific physiological process rather complex and challenging.

In late mitosis, the timely polyubiquitylation and subsequent degradation of securin and cyclin B by anaphase-promoting complex/cyclosome (APC/C) play a critical role for anaphase onset and chromosome segregation (Peters, 2006; Sullivan and Morgan, 2007). Given the essential role of APC/C in triggering chromosome segregation, it is not surprising that APC/C is a key molecular target of the spindle assembly checkpoint (SAC), which is an intricate surveillance mechanism that prolongs mitosis until all chromosomes achieve correct bipolar attachments to spindle microtubules (Murray, 2011; Musacchio, 2015). Previous studies have revealed that the active MAPK is required for spindle checkpoint activation and APC/C inhibition in *Xenopus* egg extracts or tadpole cells (Chen, 2004; Chung and Chen, 2003; Minshull et al., 1994; Takenaka et al., 1997; Zhao and Chen, 2006). These studies suggested that phosphorylation of several components of SAC, including Cdc20, Mps1 and Bub1, by MAPK serves as regulatory signals to block APC/C activation upon SAC activation (Chen, 2004; Chung and Chen, 2003; Zhao and Chen, 2006). However, all these studies used either the anti-MAPK antibodies or small molecule inhibitor UO126 to inhibit MAPK or MAPK-specific phosphatase MKP-1 to antagonize MAPK activity, these reagents could not distinguish the contributions of different MAPKs with high specificity. Thus, those seemingly defined mechanisms might suffer the ambiguity due to potential specificity issues.

The fission yeast *Schizosaccharomyces pombe* has three MAPK-signaling cascades: the pheromone signaling pathway (PSP), the stress-activated pathway (SAP) and the cell integrity pathway (CIP), with Spk1, Sty1 or Pmk1 as MAPK respectively (Figure 1A) (reviewed in (Cansado et al., 2021; Perez and Cansado, 2010)). In *S. pombe*, whether MAPKs play key roles in regulation of the APC/C and spindle checkpoint remains largely unexplored. Intriguingly, all genes encoding the major components of three MAPK-signaling cascades in fission yeast are non-essential (Kim et al., 2010), this creates the possibility for investigations on their potential functions in a genetic background with complete deletion of relevant genes.

**Figure 1.**
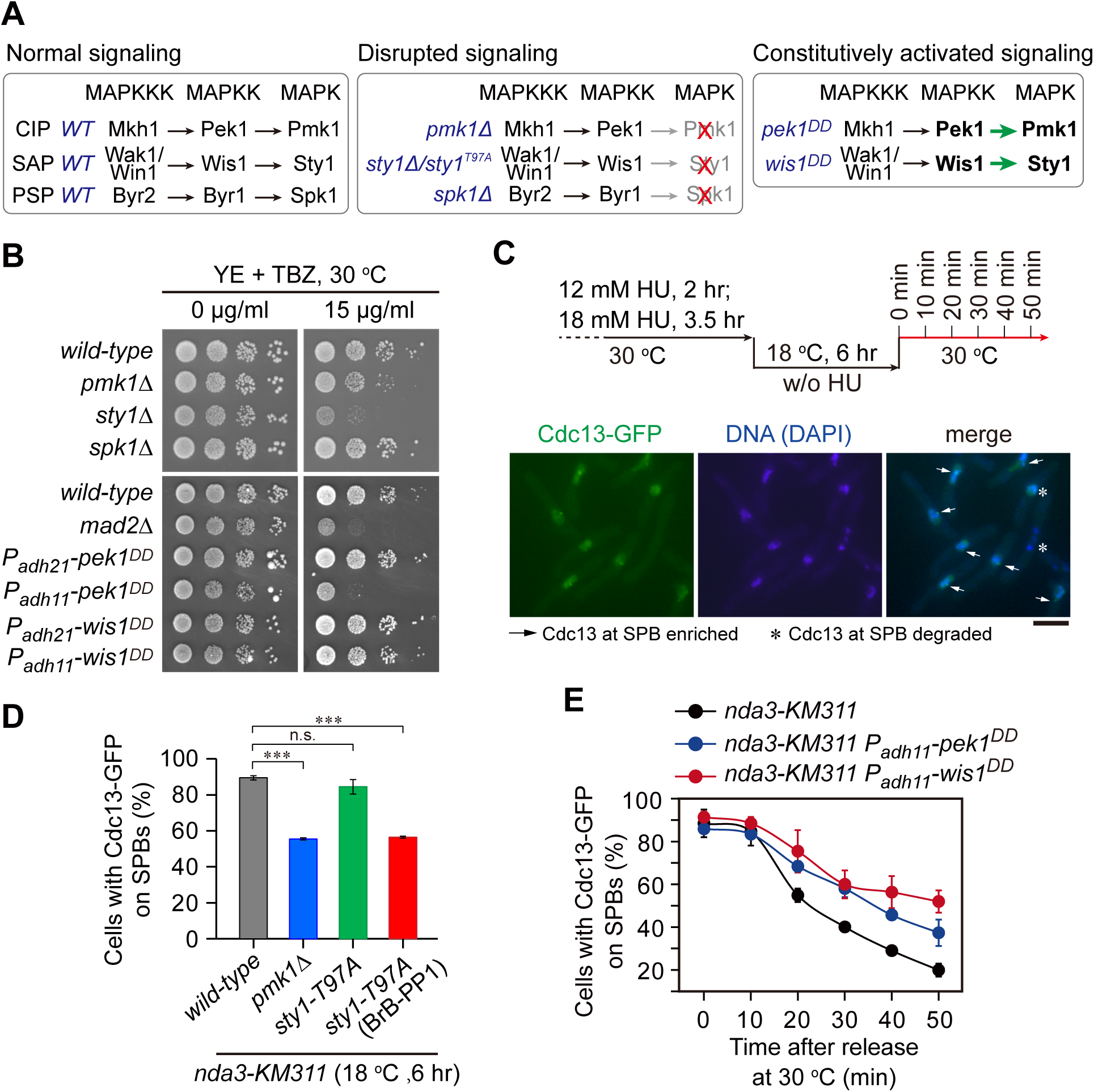
*pmk1*Δ and *sty1-T97A* mutants display spindle checkpoint defects, and *pek1*^*DD*^ and *wis1*^*DD*^ mutants display checkpoint silencing defects. (**A**) Schematic of core modules of three *Schizosaccharomyces pombe* mitogen-activated protein kinase (MAPK) signaling pathways. Each cascade consists of three core kinases (MAPKKK (MAP kinase kinase kinase), MAPKK (MAP kinase kinase), and MAPK). CIP, the cell integrity pathway; SAP, the stress-activated pathway; PSP, pheromone signaling pathway. These pathways can be disrupted by *pmk1*Δ, ATP analogue-sensitive mutation *sty1-T97A* or *spk1*Δ, and constitutively activated by mutations in MAPKKs Pek1 (*pek1*^*DD*^, *pek1-S234D;T238D*) or Wis1 (*wis1*^*DD*^, *wis1-S469D;T473D*) respectively. (**B**) Serial dilution assay on TBZ sensitivity of all MAPK deletion mutants and *pek1*^*DD*^- or *wis1*^*DD*^-overexpressing mutants. *mad2*Δ is a positive control. Note *P*_*adh11*_ is a stronger version of *P*_*adh21*_ promoter. (**C**) Schematic depiction of the experiment design for time-course analyses (in D and E) on SAC or APC/C activation. Example pictures of cells with Cdc13-GFP signals enriched or disappeared at spindle pole bodies (SPBs) are shown. Scale bar, 5 μm. (**D**) Inactivation of CIP or SAP signaling by *pmk1*Δ or *sty1-T97A* compromises full SAC activation. *sty1-T97A* was inactivated by 5μM 3-BrB-PP1. (**E**) Expression of constitutively active MAPKK *pek1*^*DD*^ or *wis1*^*DD*^ delays SAC inactivation.

In this work, we set out to directly examine the requirement of the fission yeast MAPKs in mitotic APC/C activation and anaphase entry. Our results uncovered a previously unappreciated mechanism in which two MAPKs, Pmk1 and Sty1, phosphorylate Slp1 (the Cdc20 homologue in *S. pombe*) and Mad2 respectively, upon SAC activation. Phosphorylation of Slp1^Cdc20^ promotes its degradation and subsequently impedes APC/C activation, whereas phosphorylation of Mad2 enhances inhibitory binding of mitotic checkpoint complex (Mad3-Mad2-Cdc20) to APC/C. Both phosphorylation events act in concert to sustain mitotic arrest in response to spindle defects.

## RESULTS

### SAP and CIP signaling components are required for spindle checkpoint activity

As the first step to examine the possible requirement of the fission yeast MAPKs in mitotic progression, we tested the sensitivity of deletion mutants of three fission yeast MAPKs to microtubule-destabilizing drug thiabendazole (TBZ), which has been routinely used as an indicator of defective kinetochore or spindle checkpoint (Akera et al., 2015; Saitoh et al., 1997). We noticed that *pmk1*Δ and *sty1*Δ but not *spk1*Δ cells were extremely sensitive to TBZ (Figure 1B), indicating the involvement of Pmk1 and Sty1 pathways in related processes.

To extend our analysis and precisely quantify whether the SAC is properly activated and maintained in MAPK mutants, we adopted one well-established assay using the cold-sensitive β-tubulin mutant *nda3-KM311*, which compromises the kinetochore-spindle microtubule attachment upon cold treatment (Hiraoka et al., 1984), to analyze the ability of cells to arrest in mitosis in the absence of spindle microtubules and enter anaphase when spindle microtubules are reassembled (May et al., 2017; Trautmann et al., 2004; Vanoosthuyse and Hardwick, 2009). In this assay, the SAC was first robustly activated by the *nda3-KM311* mutant at 18 °C and then inactivated simply by shift mitotically arrested cells back to permissive temperature (30 °C) (Figures 1C). Because Cdc13 (cyclin B in *S. pombe*) localizes to the spindle pole bodies (SPBs) in early mitosis and should be degraded by APC/C to promote metaphase-anaphase transition, accumulation of Cdc13-GFP on SPBs after cold treatment serves as the read-out of the SAC activation and the disappearance rate of Cdc13-GFP spot under permissive temperature reflects the SAC inactivation efficacy (Figures 1C). It is noteworthy that the ATP analogue-sensitive mutant *sty1-T97A* (i.e. *sty1-as2*) (Zuin et al., 2010), instead of *sty1*Δ deletion mutant, was used in this assay, due to the G_2_ delay in the *sty1*Δ deletion mutant (Shiozaki and Russell, 1995). We found that inactivation of CIP or SAP signaling by *pmk1*Δ or *sty1-T97A* compromised full SAC activation, as only less than 60% of cells with Cdc13-GFP on SPBs after 6 hours of cold treatment in these two mutants, whereas the percentage in wild-type after the same treatment was roughly 90% (Figure 1D and Supplemental Figure S1). Consistently, constitutively activated CIP and SAP signaling by *pek1*^*DD*^ (*pek1-S234D;T238D*) or *wis1*^*DD*^ (*wis1-S469D;T473D*) mutants (Shiozaki et al., 1998; Sugiura et al., 1999), respectively, caused yeast cells to be sensitive to TBZ and led to delayed SAC inactivation and anaphase onset after release from spindle checkpoint arrest, although the SAC activation was largely unaffected (Figure 1B, 1E and Supplemental Figure S2).

All above data suggested that MAPK pathways are evolutionarily conserved in the fission yeast to participate in spindle checkpoint activation and maintenance, of which the CIP and SAP signaling pathways, but not the Spk1 pathway, are actively involved.

### SAP and CIP signaling facilitate association of MCC with APC/C and attenuate protein levels of Slp1^Cdc20^ respectively

The mitotic checkpoint complex (MCC, consisting of Mad3-Mad2-Cdc20 in *S. pombe*) is the most potent inhibitor of the APC/C prior to anaphase (Kapanidou et al., 2017; Primorac and Musacchio, 2013; Sudakin et al., 2001). Previous study has revealed that *Xenopus* MAPK phosphorylates Cdc20 and facilitates its binding by Mad2, BubR1 (Mad3 in *Xenopus*) and Bub3 to form a tight MCC to prevent Cdc20 from activating the APC/C (Chung and Chen, 2003). In fission yeast, the key SAC components Mad2 and Mad3 and one molecule of Slp1^Cdc20^ form MCC which binds to APC/C through another molecule of Slp1^Cdc20^ upon checkpoint arrest, and the recovery from mitotic arrest accompanies the loss of MCC-APC/C binding (May et al., 2017; Sczaniecka et al., 2008; Sewart and Hauf, 2017; Vanoosthuyse and Hardwick, 2009). Since we found that constitutive activation of CIP or SAP signaling in fission yeast by *pek1*^*DD*^ or *wis1*^*DD*^ mutations caused prolonged SAC activation, we were suspicious that these two MAPK pathways might also execute their functions cooperatively through a similar mechanism.

To test this possibility, we analyzed the MCC occupancy on APC/C by immunoprecipitations of the APC/C subunit Lid1 (Yoon et al., 2002) in *pek1*^*DD*^ and *wis1*^*DD*^ cells arrested by activated checkpoint. Compared to wild-type, more MCC components (Mad2+Mad3 and Slp1^Cdc20^) were co-immunoprecipitated in *wis1*^*DD*^ cells (Figure 2A), while less of them were co-immunoprecipitated in *pek1*^*DD*^ cells (Supplemental Figure S3A), suggesting that activation of the SAP but not the CIP pathway enhanced the association of MCC with APC/C. The significantly decrease of co-immunoprecipitated Mad2, Mad3 and Slp1^Cdc20^ in *sty1-T97A* mutant further supported this conclusion (Figure 2A). We also examined the MCC assembly by immunoprecipitations of Mad3 in *wis1*^*DD*^ and *sty1-T97A* cells, and found that the amount of associated Mad2 and Slp1 was not altered (Supplemental Figure S3B), indicating that SAP signaling did not affect the assembly of MCC.

**Figure 2.**
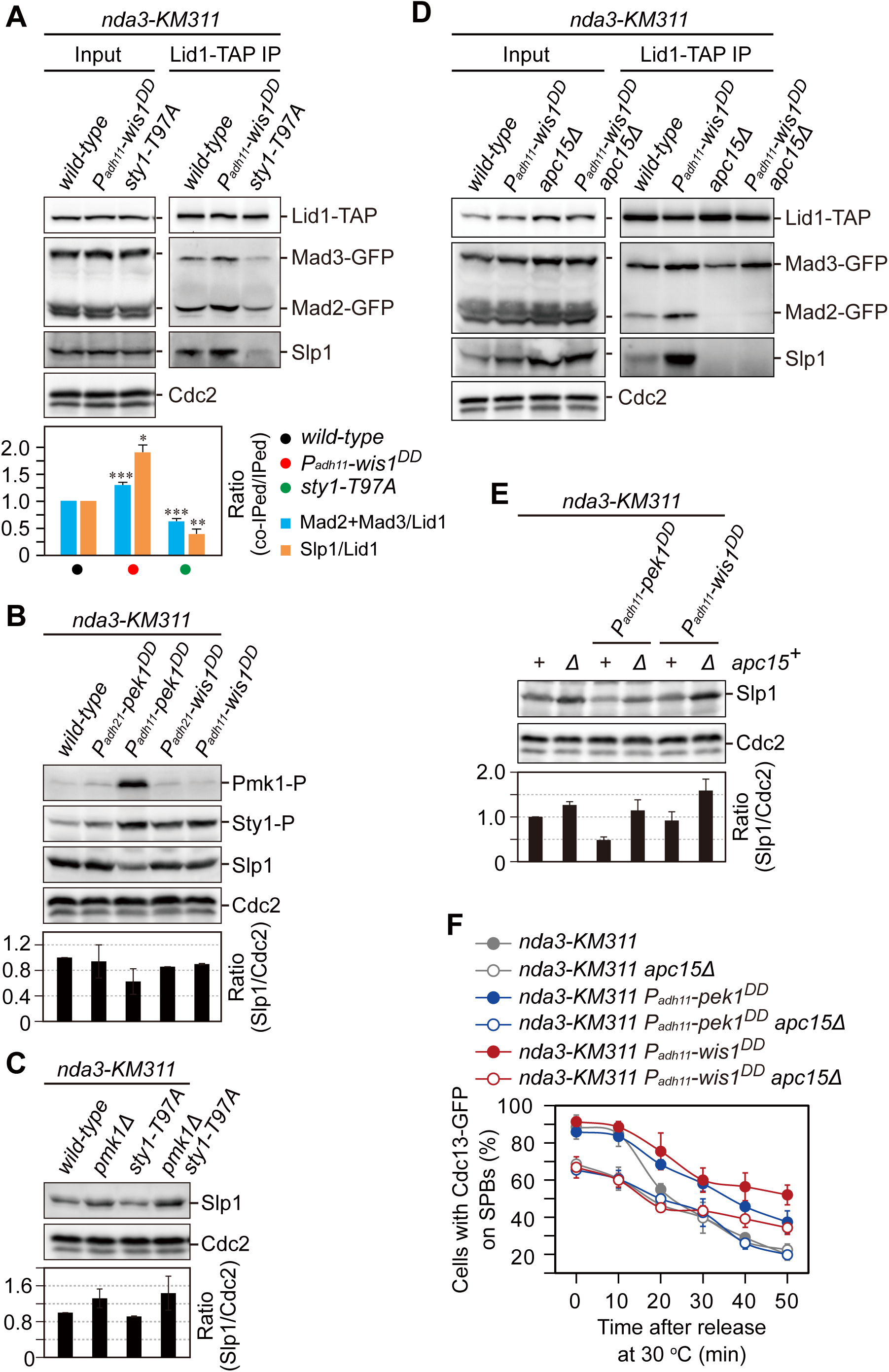
Upon SAC activation, SAP and CIP signaling facilitate association of MCC with APC/C and attenuate protein level of Slp1^Cdc20^ respectively. (**A**) Upon SAC activation, constitutive activation of Wis1 (*wis1*^*DD*^) enhances and removal of Sty1 (*sty1-T97A*) reduces the association of MCC with APC/C. The association of Mad2, Mad3 and Slp1^Cdc20^ to Lid1 was assessed by immunoprecipitation of Lid1-TAP in checkpoint-arrested cells. The amount of co-immunoprecipitated Mad2, Mad3 and Slp1 was normalized to those of total immunoprecipitated Lid1 in each sample, with the relative ratio between Mad2-GFP plus Mad3-GFP or Slp1 and Lid1-TAP in wild-type sample set as 1.0. Note that more Mad2, Mad3 and Slp1^Cdc20^ was co-immunoprecipitated in *wis1*^*DD*^ cells compared to wild-type cells. *sty1-T97A* was inactivated by 5μM 3-BrB-PP1. (**B, C**) Upon SAC activation, Slp1^Cdc20^ levels are increased in *pmk1*Δ cells and reduced in *pek1*^*DD*^ compared to wild-type cells, and Slp1^Cdc20^ abundance remained unaltered in *sty1-T97A* and *wis1*^*DD*^ mutants. Slp1^Cdc20^ levels were quantified with the relative ratio between Slp1^Cdc20^ and Cdc2 in wild-type strain set as 1.0. *sty1-T97A* was inactivated by 5μM 3-BrB-PP1. Phosphorylated Pmk1 (Pmk1-P) or phosphorylated Sty1 (Sty1-P) represents activated CIP or SAP signaling, respectively. The experiments were repeated 3 times and the mean value for each sample was calculated. (**D**) *apc15*Δ abolishes the elevated binding of MCC to APC/C in *wis1*^*DD*^ cells. (**E**) *apc15*Δ restores Slp1 levels in *pek1*^*DD*^ mutant. (**F**) *apc15*Δ relieves delayed APC/C activation and rescues SAC inactivation defects in both *pek1*^*DD*^ and *wis1*^*DD*^ mutants.

The decrease of co-immunoprecipitated Mad2 and Mad3 in *pek1*^*DD*^ cells was largely out of expectation. Further analysis of the inputs of these immunoprecipitations showed that Slp1^Cdc20^ levels were significantly reduced in *pek1*^*DD*^ but not *wis1*^*DD*^ cells (Supplemental Figure S3A). Our immunoblotting analyses using the total lysate of *pek1*^*DD*^ or *wis1*^*DD*^ cells arrested at metaphase also confirmed these results (Figure 2B). Consistently, the Slp1^Cdc20^ level was significantly increased in *pmk1*Δ and *pmk1 Δ sty1-T79A* cells but not in *sty1-T79A* cells (Figure 2C). These results collectively revealed a negative effect of CIP but not SAP signaling on Slp1^Cdc20^ levels, which further affects the MCC-APC/C interaction in CIP mutants.

The increased Slp1^Cdc20^ levels in *pmk1*Δ cells is reminiscent of the previous observations in *apc15*Δ mutant, which also shows elevated levels of Slp1^Cdc20^ (May et al., 2017; Sewart and Hauf, 2017). It has been shown in both human and fission yeast cells that Apc15 mediates MCC binding to APC/C and is required for Cdc20/Slp1 autoubiquitylation and its turnover by APC/C (Mansfeld et al., 2011; May et al., 2017; Sewart and Hauf, 2017; Uzunova et al., 2012). We reasoned that the absence of Apc15 may reverse the negative effect of *pek1*^*DD*^ on Slp1^Cdc20^ levels and the positive effect of *wis1*^*DD*^ on MCC-APC/C association. As expected, we indeed observed that deletion of *apc15* relieved the enhanced MCC-APC/C interaction in *wis1*^*DD*^ cells and recovered Slp1^Cdc20^ levels in *pek1*^*DD*^ cells (Figure 2D and 2E), indicating that attenuation of Slp1^Cdc20^ levels by Pek1^DD^ is very likely through facilitating its autoubiquitylation. Furthermore, the absence of Apc15 also lowered the spindle assembly checkpoint response to disruption of spindles and accelerated anaphase entry in both *pek1*^*DD*^ and *wis1*^*DD*^ cells (Figure 2F). These results are also consistent with previous report showing the relative abundance between checkpoint proteins and the checkpoint target Slp1 is an important determinant of checkpoint robustness (Heinrich et al.,2013).

Together, our above data suggested a new mechanism involves division of labor between the two fission yeast MAPK pathways SAP (Wis1-Sty1) and CIP (Pek1-Pmk1) upon SAC activation, which is required in concert to enhance MCC affinity for APC/C and lower Slp1^Cdc20^ levels to strongly inhibit APC/C activity.

### Sty1 enhances MCC-APC/C interaction through phosphorylation of Mad2

Our observation that the constitutively activated SAP signaling by *wis1*^*DD*^ mutation enhanced association of MCC with APC/C (Figures 2A, 2C and Supplemental Figure S3) prompted us to examine the possibility that the SAP signaling pathway is directly involved in regulating this interaction.

First, an *in vitro* phosphorylation reaction was performed, and we found that, in the presence of *wis1*^*DD*^ and functional Sty1 (but not the *sty1-T79A* mutant with the addition of 3-BrB-PP1), Mad2 but not Mad3 could be phosphorylated (Supplemental Figure S4 A and C), suggesting activated Sty1 can phosphorylate Mad2 *in vitro*. The samples from these assays were further analyzed by mass-spectrometry and 13 *in vitro* Sty1 sites in Mad2 were identified (Figure 3A).

**Figure 3.**
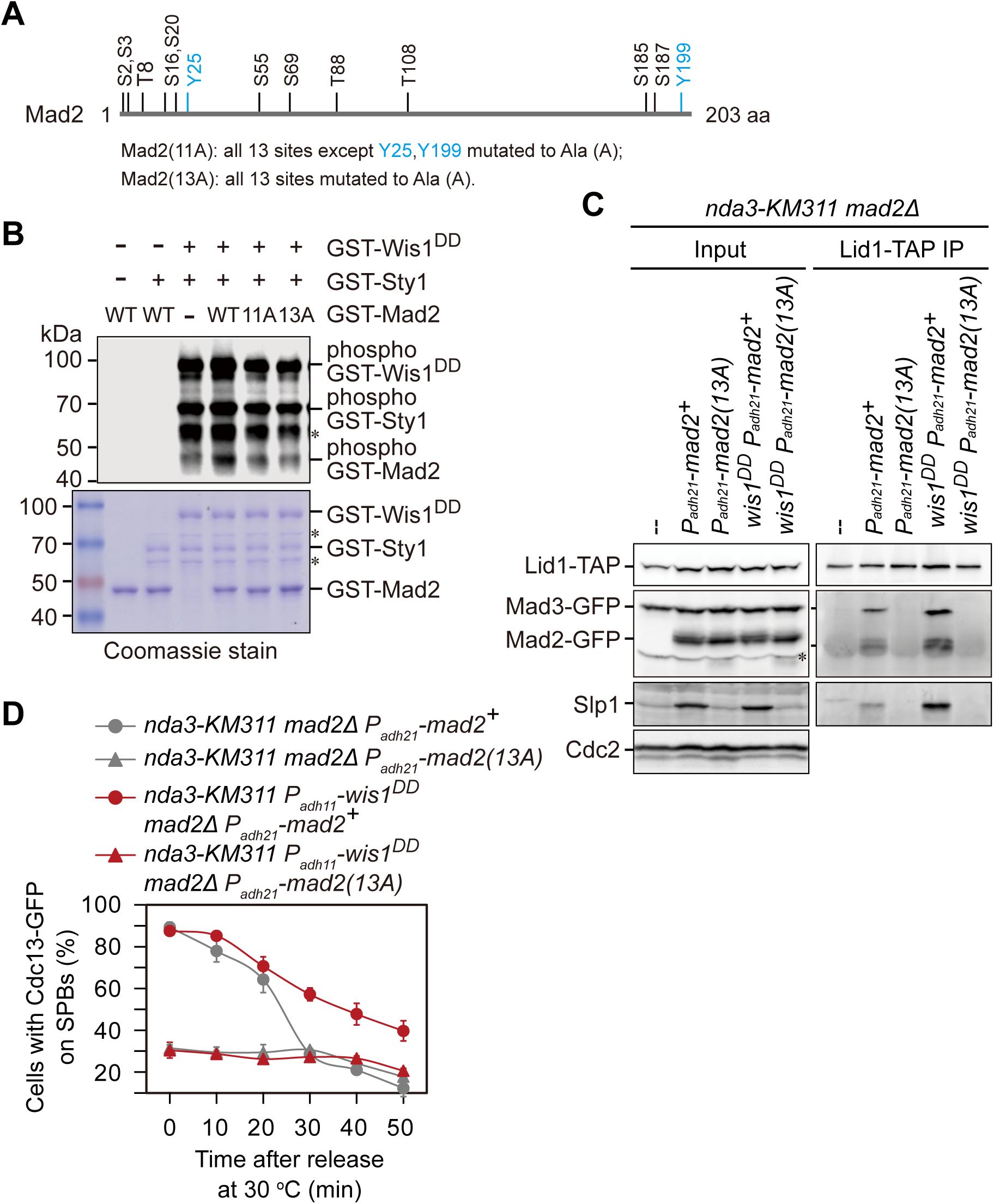
Sty1 enhances spindle checkpoint activity through phosphorylation of Mad2. (**A**) Schematic depiction of the Mad2 protein with the substitutions of the 11 or 13 putative Sty1 phosphorylation sites to alanines (11A/13A) indicated. (**B**) Sty1 phosphorylates Mad2 *in vitro*. Representative results of non-radioactive kinase assay using recombinant GST-fused Mad2, Mad2(11A) or Mad2(13A) and Sty1 and Wis1^DD^ are shown. (**C**) *mad2*^*13A*^ mutation abolishes the enhanced MCC-APC/C association in *wis1*^*DD*^ mutant. Note that more Mad2, Mad3 and Slp1^Cdc20^ were co-immunoprecipitated in *wis1*^*DD*^ cells compared to wild-type cells when wild-type but not mutant Mad2 was present. (**D**) Delayed APC/C activation and SAC inactivation in *wis1*^*DD*^ mutant are reversed by *mad2*^*13A*^ mutation.

Next, through multiple sequence alignment analysis, we found that some of these identified phosphorylation sites in Mad2 (*e*.*g*., Ser55, Ser185 and Ser187) are highly conserved in a few fungi species and humans (Supplemental Figure S5). However, mutations of these sites had no disruptive effect on either MCC-APC/C interaction or activation and inactivation of SAC, as the amount of co-immunoprecipitated MCC components was not altered in these mutants (Supplemental Figure S6A), and the dynamics of Cdc13-GFP on SPBs after release from cold treatment were actually the same in these mutants as in wild-type (Supplemental Figure S6B). These results suggested that elevated MCC affinity to APC/C promoted by Sty1 requires multi-site phosphorylation of Mad2.

Finally, we substituted all 13 potential phosphorylation sites by alanine (*mad2*^*13A*^), and found that this mutant abolished the enhanced association of MCC with APC/C (Figures 3C) and also rescued APC/C activation defects in *wis1*^*DD*^ mutant (and 3D).

Together, these data suggested that the SAP signaling promotes MCC-APC/C interaction through phosphorylating MCC component Mad2. This mechanism is distinct from that in previous report demonstrating instead phosphorylation of Cdc20 by MAPK is responsible for its facilitated association with MCC to prevent Cdc20 from activating the APC/C in *Xenopus* egg extracts and tadpole cells (Chung and Chen, 2003).

### Pmk1 restrains APC/C activity through binding to and phosphorylation of Slp1 upon SAC activation

To test whether two MAPKs Pmk1 and Sty1 possibly phosphorylate Slp1^Cdc20^, we first examined the potential interaction of MAPKs with Slp1^Cdc20^. We found that only Pmk1 but not Sty1 directly bound to Slp1^Cdc20^, though both kinases could be pull-downed from yeast cell lysates by Slp1^Cdc20^ (Supplemental Figure S7 and data not shown).

Next, we attempted to find the key residues in Slp1^Cdc20^ that mediate the Pmk1-Slp1^Cdc20^ interaction. In previous reports, MAPK-docking sites, which are important for interaction with MAPKs, have been identified at the N termini of different MAPKKs from yeast to humans with a notable feature of a cluster of at least two basic residues (mainly Lys and Arg) separated by a spacer of 2-6 residues from a hydrophobic-X-hydrophobic sequence (Bardwell et al., 2001; Bardwell and Thorner, 1996). By visual scanning of Slp1^Cdc20^ sequence, we noticed five basic-residue patches which loosely resemble the consensus sequence of the known MAPK-docking sites, and four of them are within N-terminal portion of the protein (Supplemental Figure S8). However, only mutating (Lys/Arg to Glu) either of the two most N-terminal basic-residue patches (19-KKR-21 and 47-KR-48) in Slp1^Cdc20^ resulted in decreased amount of Slp1^Cdc20^ pulled down by Pmk1-GST *in vitro* (Supplemental Figure S7B), and mutating both completely abolished their interaction (Figure 4A and 4B), demonstrating that these two short basic patches serve as the major docking sites for Pmk1. Importantly, mutations of these basic residues to glutamate (E) mimicked *pmk1*Δ mutant in both elevated Slp1^Cdc20^ protein levels (Figure 4C) and defective activation and maintenance of the SAC upon *nda3*-mediated checkpoint arrest (Figure 4D and Supplemental Figure S9).

**Figure 4.**
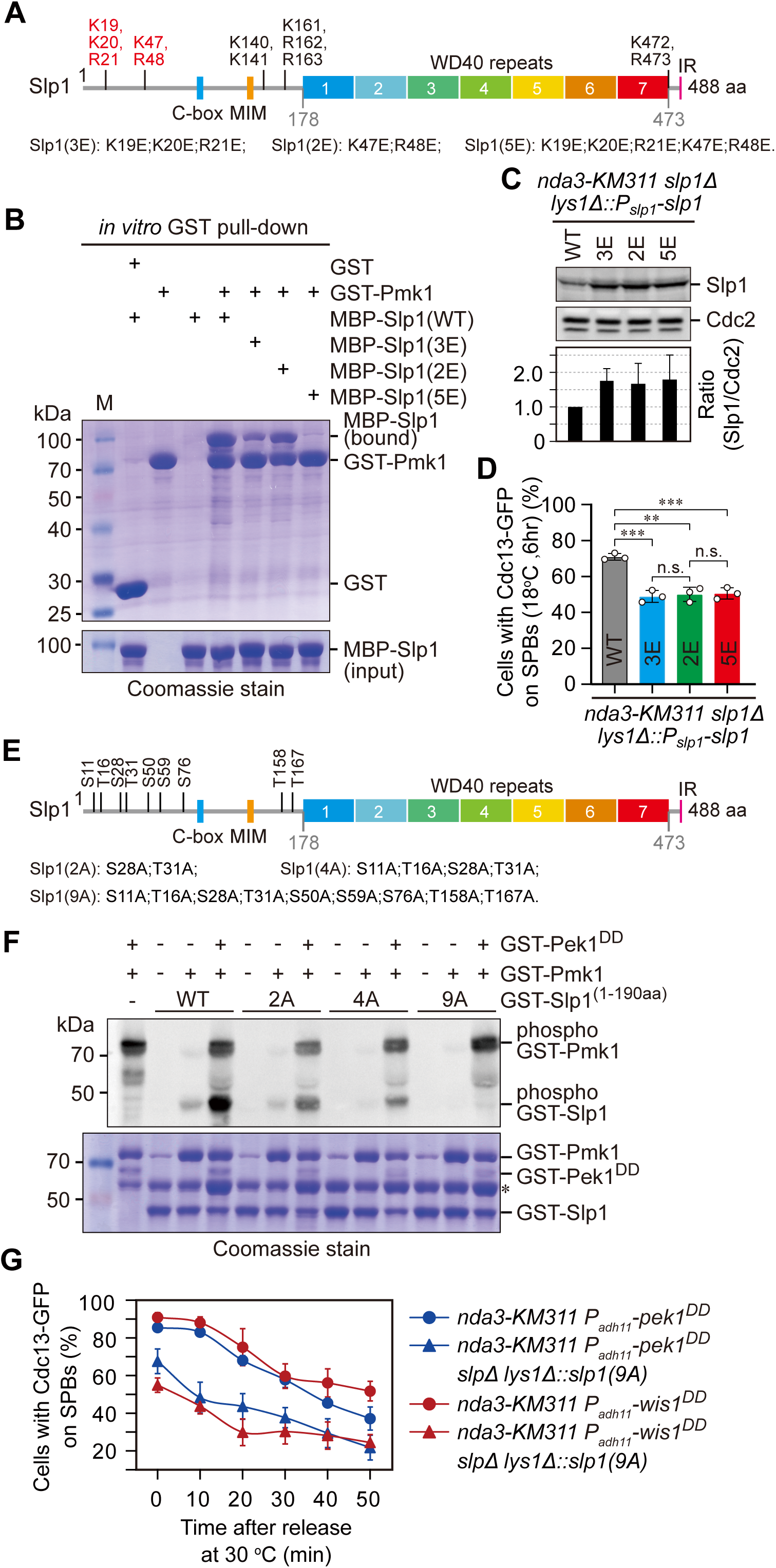
Pmk1 binds and phosphorylates Slp1^Cdc20^ to attenuate its levels upon SAC activation. (**A**) Schematic depiction of the *S. pombe* Slp1 protein with five potential basic-residue patches required for Slp1-Pmk1 association indicated, and two confirmed basic-residue patches highlighted in red. MIM, Mad2-interaction motif; IR, isoleucine-arginine tail. (**B**) Two mini basic-residue patches (19-KKR-21 and 47-KR-48) within N-terminus of Slp1 mediate the direct interaction between Slp1 and Pmk1. *In vitro* GST pull-down assays were performed with bacterially expressed recombinant proteins. (**C**) Mutation of two basic-residue patches of Slp1 stabilizes Slp1 protein upon SAC activation. Slp1^Cdc20^ levels were quantified with the relative ratio between Slp1^Cdc20^ and Cdc2 in wild-type strain set as 1.0. The experiments were repeated 3 times and the mean value for each sample was calculated. (**D**) Mutation of two basic-residue patches of Slp1 compromises SAC full activation and maintenance. Cells were first grown at 30 °C and then arrested at 18 °C for 6 hours. The percentage of cells with Cdc13-GFP on SPBs was assessed. (**E**) Schematic depiction of Slp1 with nine putative Pmk1 phosphorylation sites indicated. (**F**) Slp1 phosphorylation by Pmk1 was analyzed by the *in vitro* non-radioactive kinase assay using recombinant GST-fused Slp1 fragment 1-190 aa harboring wild-type or alanine (A) substitutions (2A, 4A or 9A) and full-length Pmk1 and Pek1^DD^. Asterisk indicates the unspecific proteins after Coomassie blue staining. Results are representative of three independent experiments. (**G**) Delayed APC/C activation and SAC inactivation in *pek1*^*DD*^ and *wis1*^*DD*^ mutants are reversed by *slp1*^*9A*^ mutation.

Further, by motif search and visual inspection, we found nine potential MAPK phosphorylation consensuses (SP/TP) present within the N-terminal sequence of Slp1^Cdc20^ (Figures 4E and Supplemental Figure S8B). Strikingly, Pmk1 could sufficiently phosphorylate Slp1^Cdc20^ *in vitro* with the presence of *pek1*^*DD*^ (Figure 4F). In the same reaction setup, the levels of phosphorylated Slp1^Cdc20^ harboring 2 or 4 Ala substitutions (Slp1^2A^ or Slp1^4A^) for Ser or Thr of potential MAPK phosphorylation sites were significantly decreased, and Slp1^9A^ was completely non-phosphorylatable by Pmk1 (Figure 4F), suggesting these sites are all potential Pmk1 targets. Among nine putative MAPK phosphorylation sites, S28 and T31 are most likely the major contributing sites for influencing APC/C activation, since the *slp1*^*S28A;T31A*^ mutant has similar percentage of Cdc13-GFP on SPBs after cold treatment to the *slp1*^*9A*^ mutant (Supplemental Figure S10). These sites also show best fit with the stringent MAPK phosphorylation consensus P-X-S/TP (Supplemental Figure S8B).

Together, these findings indicate that the CIP signaling restrains APC/C activity through direct binding and phosphorylating Slp1^Cdc20^ to lower its levels, likely by facilitating autoubiquitylation of phosphorylated Slp1^Cdc20^.

## DISCUSSION

So far, the only known cell-cycle control stage linked to MAPKs in fission yeast is at G_2_/M transition, where the p38 MAPK family member Sty1 either arrests or promotes mitotic commitment in unperturbed, stressed or resumed cell cycles after stress depending on the downstream kinases it associates with. Sty1 can phosphorylate kinase Srk1 at threonine 463 (T463), which further phosphorylates and inhibits Cdc25 phosphatase to negatively regulate the onset of mitosis (Lopez-Aviles et al., 2005; Lopez-Aviles et al., 2008; Smith et al., 2002). Sty1 also mediates phosphorylation of Polo kinase Plo1 at serine 402 (S402), which then promotes its recruitment to the spindle pole bodies (SPBs), where it positively modulates Cdc25 control of Cdc2 (fission yeast Cdk1) activity for mitotic entry (Petersen and Hagan, 2005; Petersen and Nurse, 2007).

In the current study, we have expanded MAPK signaling-controlled cell-cycle stage to metaphase-anaphase transition in fission yeast. In response to unattached or tensionless kinetochores, the spindle assembly checkpoint (SAC) generates the mitotic checkpoint complex (MCC), which inhibits the APC/C and block anaphase onset by suppressing APC/C-catalyzed ubiquitination and subsequent degradation of securin and cyclin B. As a critical APC/C inhibitor, MCC fulfills its function through the physical binding of BubR1/Mad3 and Mad2 to two molecules of Cdc20/Slp1 (i.e. Cdc20^MCC^ and Cdc20^APC/C^), this mechanism has been confirmed at least in both humans and fission yeast (Alfieri et al., 2016; Chao et al., 2012; Izawa and Pines, 2015; Sewart and Hauf, 2017; Yamaguchi et al., 2016). In this study, we have identified novel phosphorylation events of Mad2 and Slp1^Cdc20^ by two fission yeast MAPKs from respective SAP and CIP signaling pathways upon SAC activation, and thus established another critical APC/C-inhibitory mechanism in the spindle checkpoint. In this newly uncovered dual mechanism, two MAPKs restrain APC/C activity through distinct manners, either by phosphorylating Slp1^Cdc20^ to lower its levels or by phosphorylating Mad2 to promote MCC-APC/C association, respectively (Figure 5A). The regulation of mitotic entry by MAPK in fission yeast is triggered by many environmental insults, including high osmolarity, oxidative stress, heat shock, centrifugation, nutrient starvation and rapamycin treatment (Hartmuth and Petersen, 2009; Petersen and Hagan, 2005; Petersen and Nurse, 2007; Shiozaki and Russell, 1995; Shiozaki et al., 1998), the extracellular or intracellular cue sensed by MAPKs during anaphase entry remains to be elucidated.

**Figure 5.**
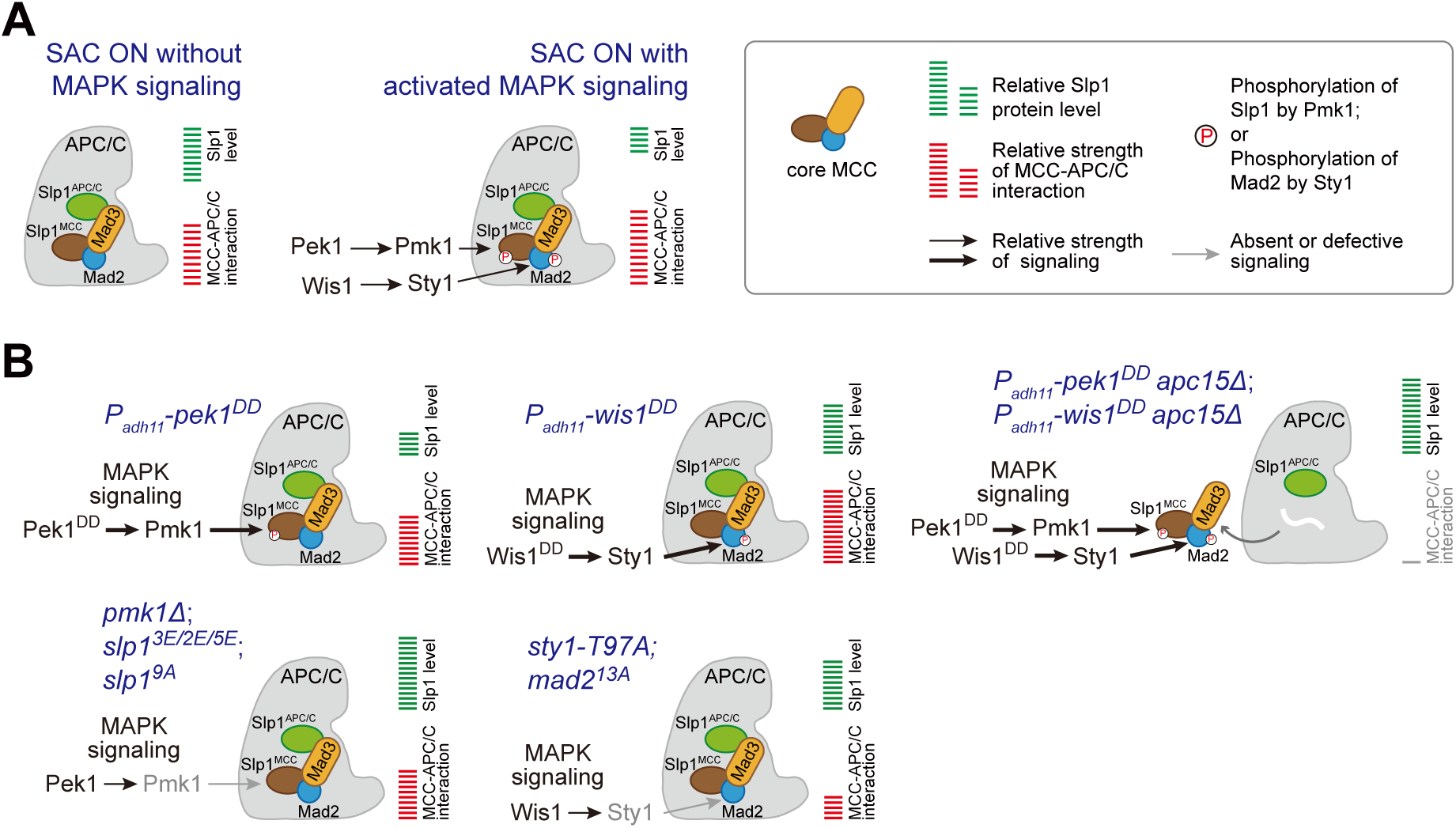
Summary of the mechanisms that how MAPKs negatively regulate APC/C activity in fission yeast. (**A**) Schematic depiction of possible mechanisms that how activated MAPK signaling pathways are involved in delaying APC/C activation and SAC inactivation. Upon SAC and MAPK signaling activation, division of labor between the CIP and the SAP pathways enables the phosphorylation of Slp1 and Mad2 by Pmk1 and Sty1 respectively, which leads to lowered Slp1 level and enhanced MCC affinity for APC/C. (**B**) Summary of the consequences of constitutively activated CIP or SAP signaling (Pek1^DD^ or Wis1^DD^), and mutations of *pmk1 Δ, sty1-T97A, slp1*^*9A*^, *mad2*^*13A*^ or *apc15*Δ on Slp1^Cdc20^ protein levels and MCC-APC/C association. White wavy line indicates the absence of Apc15.

In previous studies, two fission yeast MCC components, Mad2 and Mad3, have been demonstrated to be phosphorylated by Mph1^Mps1^, the master kinase for spindle checkpoint activation (Zich et al., 2016; Zich et al., 2012). These posttranslational modifications of the MCC are required to stabilize the MCC-APC/C interaction and thereby inhibit APC/C activity during checkpoint arrest (Zich et al., 2016; Zich et al., 2012). Interestingly, we found that activated SAP signaling also similarly promotes the MCC-APC/C interaction. We identified 13 *in vitro* Sty1-phosphorylated residues that are scattered in fission yeast Mad2. Although some of them, such as Ser55, Ser185 and Ser187, are also highly conserved in humans (Supplemental Figure S5), our mutation analyses suggested that phosphorylation of multiple residues in addition to these sites in Mad2 is required for Sty1-mediated effect on MCC-APC/C affinity (Supplemental Figure S6). It will be interesting to address whether this MAPK-mediated Mad2 phosphorylation exist as a general mechanism for fostering spindle checkpoint activity in higher eukaryotes. However, none of the 11 serine or threonine residues we identified in fission yeast Mad2 fits the canonical MAPK phosphorylation motif, which requires a proline at the P+1 position and frequently with proline at the P-2 position of serine or threonine residues (Gonzalez et al., 1991; Manke et al., 2005). It is possible that the potential MAPK phosphorylation sites in metazoan Mad2 have been overlooked by searching for only the optimal substrate phosphorylation motif. Actually, several confirmed MAPK phosphorylation substrates indeed contain non-canonical sites, for example, the serine 402 (S402) in fission yeast Plo1 phosphorylated by Sty1 (Petersen and Hagan, 2005), the serines 309 and 361 (S309 and S361) in human Cdc25B and serine 216 (S216) in human Cdc25C phosphorylated by p38 (Bulavin et al., 2001).

APC/C activity can be regulated at multiple levels, Cdk1-dependent APC/C phosphorylation is one of the major prerequisites for APC/C activation and anaphase onset, in which Apc3 and Apc1 are key subunits for positive phospho-regulation (Fujimitsu et al., 2016; Kraft et al., 2003; Qiao et al., 2016; Steen et al., 2008; Zhang et al., 2016). However, Cdk1-dependent phosphorylation of APC/C co-activator Cdc20 lessens its interaction with APC/C, thus serves as a negative phospho-regulation (Labit et al., 2012). In addition to its phosphorylation by Cdk1, phosphorylation of Cdc20 by multiple other kinases, such as Bub1, PKA, MAPK and Plk1, has also been demonstrated in *Xenopus*, humans or budding yeast to be inhibitory to APC/C activation and improper anaphase onset upon SAC activation ((Jia et al., 2016) and reviewed in (Yu, 2007)). Therefore, Cdc20 serves as an integrator of multiple intracellular signaling cascades that regulate progression through mitosis.

The current model of inhibitory signaling from Cdc20 posits that its phosphorylation either promotes the formation of MCC or inhibits APC/C^Cdc20^ catalytically, both are required for proper spindle checkpoint signaling. Interestingly, our current study establishes Slp1^Cdc20^ phosphorylation by fission yeast MAPK Pmk1 as another critical APC/C-inhibitory mechanism in the spindle checkpoint through reducing the protein levels of Slp1^Cdc20^. Actually, previous studies in fission and budding yeast have underlined the importance of accurate relative abundance between checkpoint proteins and Cdc20/Slp1^Cdc20^, which sets an important determinant of checkpoint robustness (Heinrich et al., 2013; Pan and Chen, 2004). A benefit of allowing the association of MCC-bound Cdc20 with APC/C is to provide an opportunity for Cdc20 autoubiquitylation by APC/C, a process aided by APC/C subunit Apc15 (Mansfeld et al., 2011; May et al., 2017; Sewart and Hauf, 2017; Uzunova et al., 2012). Notably, the depletion of *apc15*^*+*^ restores Slp1^Cdc20^ levels in *pek1*^*DD*^ cells (Figure 5B), consistent with the idea that Pmk1-mediated Slp1^Cdc20^ phosphorylation very likely facilitates its autoubiquitylation.

In summary, our studies reveal a distinct dual mechanism of MAPK-dependent anaphase onset delay imposed directly on APC/C co-activator Slp1^Cdc20^ and the key component of MCC Mad2, which may emerge as an important regulatory layer to fine tune APC/C and MCC activity upon SAC activation. Importantly, although this mechanism of APC/C regulation by two MAPKs has not been examined in other organisms, it is very likely evolutionarily conserved in higher eukaryotes including humans, given the involvement of MAPKs in general response to wide range of adverse situations and similar regulations of APC/C and SAC in eukaryotes characterized so far. It is also plausible to assume that the MAPK-mediated APC/C inhibition pathway may also contribute to the pathogenesis of multiple cancers.

## METHODS

### Fission yeast strains, media and genetic methods

Fission yeast cells were grown in either YE (yeast extract) rich medium or EMM (Edinburgh minimal medium) containing the necessary supplements. G418 disulfate (Sigma-Aldrich; A1720), hygromycin B (Sangon Biotech; A600230) or nourseothiricin (clonNAT; Werner BioAgents; CAS#96736-11-7) was used at a final concentration of 100 μg/ml and thiabendazole (TBZ) (Sigma-Aldrich; T8904) at 5-15 μg/ml in YE media where appropriate. For serial dilution spot assays, 10-fold dilutions of a mid-log-phase culture were plated on the indicated media and grown for 3 to 5 days at indicated temperatures. To create strain with an ectopic copy of *slp1*^*+*^ at *lys1*^*+*^ locus on chromosome 1, the open reading frame of *slp1*^*+*^ (1467 bp) and its upstream 1504 bp sequence was first cloned into the vector pUC119-*P*_*adh21*_-MCS-*hphMX6-lys1*\* with *adh21* promoter (*P*_*adh21*_) removed using the ‘T-type’ enzyme-free cloning method (Chen et al., 2017). The resultant plasmid was linearized by *Apa*I and integrated into the *lys1*^*+*^ locus, generating the strain *lys1Δ::P*_*slp1*_*-slp1*^*+*^*-T*_*adh1*_*::hphMX6*. To generate the *slp1(2A)/(4A)/(9A)* strains, only two sites (Ser28 and Thr31), the first four (Ser11, Thr16, Ser28 and Thr31) or all 9 sites of *slp1*^*+*^ (Ser11, Thr16, Ser28, Thr31, Ser50, Ser59, Ser76, Thr158 and Thr167) in pUC119-*P*_*slp1*_*-slp1*^*+*^*-T*_*adh1*_*::hphMX6* were changed to alanines (A) using standard methods of site-directed mutagenesis. The resulting plasmids were linearized and integrated into the *lys1*^*+*^ locus of chromosome 1 using the *hyg*^*r*^ marker. Then endogenous *slp1*^*+*^ was deleted and replaced with *ura4*^*+*^. To generate the constitutively active MAPKK strains *pek1*^*DD*^ or *wis1*^*DD*^, the Ser234 and Thr238 of *pek1*^*+*^ in pUC119-*P*_*adh21*_-*6HA-pek1*^*+*^-*T*_*adh1*_-*hphMX6-lys1*\*, pUC119-*P*_*adh11*_-*6HA-pek1*^*+*^-*T*_*adh1*_-*hphMX6-lys1*\* and pUC119-*P*_*adh11*_-*6HA-pek1*^*+*^-*T*_*adh1*_-*kanMX6-ura4*^*+*^ and the Ser469 and Thr473 of *wis1*^*+*^ in pUC119-*P*_*adh21*_-*6HA-wis1*^*+*^-*T*_*adh1*_-*hphMX6-lys1*\*, pUC119-*P*_*adh11*_-*6HA-wis1*^*+*^-*T*_*adh1*_-*hphMX6-lys1*\* and pUC119-*P*_*adh11*_-*6HA-wis1*^*+*^-*T*_*adh1*_-*kanMX6-ura4*^*+*^ were changed to aspartic acids (D) by standard methods of site-directed mutagenesis. To generate the *mad2(1A)/(2A)/(3A)/(11A)/(13A)* strains, individual or all 13 sites of *mad2*^*+*^ (Ser2, Ser3, Thr8, Ser16, Ser20, Tyr25, Ser55, Ser69, Thr88, Thr108, Ser185, Ser187, Tyr199) in pUC119-*P*_*adh21*_*-mad2*^*+*^*-T*_*adh1*_*::hphMX6*, pUC119-*P*_*adh21*_*-mad2*^*+*^*-GFP-T*_*adh1*_*::natMX6* or pUC119-*P*_*mad2*_*-mad2*^*+*^*-T*_*adh1*_*::natMX6* were changed to alanines (A) using standard methods of site-directed mutagenesis. The above plasmids were then linearized and integrated into the *lys1*^*+*^ locus of chromosome 1 using the *hyg*^*R*^ marker, the *ura4*^*+*^ locus on chromosome 3 using the *kan*^*R*^ marker, or the *ade6*^*+*^ locus on chromosome 3 using the *nat*^*R*^ marker, respectively. A list of the yeast strains used is in Supplemental Table S1.

### Fission yeast cell synchronization methods

For *nda3-KM311* strains, cells were grown at the permissive temperature for *nda3-KM311* (30 °C) to mid -log phase, synchronized at S phase by adding HU (Sangon Biotech; A600528) to a final concentration of 12 mM for 2 hours followed by a second dose of HU (6 mM final concentration) for 3.5 hours. HU was then washed out and cells were released at specific temperatures as required by subsequent experiments.

### Spindle checkpoint activation and silencing assays in fission yeast

For checkpoint silencing assay in the absence of microtubules, mid-log *cdc13-GFP nda3-KM311* cells were first synchronized with HU at 30 °C and then arrested in early mitosis by shifting to 18 °C for 6 hr, followed by incubation at 30 °C to allow spindle reformation and therefore spindle checkpoint inactivation. For strains carrying *sty1-T97A*, cells were treated with 5μM ATP analog 3-BrB-PP1 (Toronto Research Chemicals; A602985) during 18 °C incubation for 1.5 hr to inactivate Sty1 kinase activity before being collected. Cells were withdrawn at certain time intervals and fixed with cold methanol and stained with DAPI. 200-300 cells were analyzed for each time point. Each experiment was repeated at least three times.

### Immunoblotting and immunoprecipitation

For Western blot and immunoprecipitation experiments, yeast cells were collected from *nda3-KM311* arrested and released cultures, followed by lysing with glass bead disruption using FastPrep24 homogenizer (MP Biomedical) in NP40 lysis buffer (6 mM Na_2_HPO_4_, 4 mM NaH_2_PO4, 1% NP-40, 150 mM NaCl, 2 mM EDTA, 50 mM NaF, 0.1 mM Na_3_VO_4_) plus protease inhibitors as previously described (Wang et al., 2012). Proteins were immunoprecipitated by IgG Sepharose beads (GE Healthcare; 17-0969-01) (for Lid1-TAP) or GFP-Trap beads (ChromoTek) (for Mad3-GFP). Immunoblot analysis of cell lysates and immunoprecipitates was performed using appropriate antibodies at 1:500 - 1:5000 dilutions and was read out using chemiluminescence.

### Expression and purification of recombinant proteins

6His, GST or MBP fusion constructs of fission yeast Slp1^Cdc20(full-length)^, Slp1^Cdc20(1-190aa)^, Pmk1, Sty1, Pek1, Wis1, Mad2, Mad3 and Atf1 were generated by PCR of the corresponding gene fragments from yeast genomic DNA or cDNA and cloned in frame into expression vectors pET28a(+) (Novagen), pGEX-4T-1 (GE Healthcare) or pMAL-c2x (New England Biolabs), respectively. Site-directed mutagenesis was done by standard methods to generate vectors carrying Pek1-DD(S234D, T238D), Wis1-DD(S469D, T473D), Slp1^Cdc20^(2A), Slp1^Cdc20^(4A), Slp1^Cdc20^(9A), Mad2(11A), Mad2(13A), or Pmk1 carrying K(Lys)/R(Arg) to E(Glu) mutations at individual or combined sites including K19, K20, R21, K47, R48, K140, K141, K161, R162, R163, K472, R473. Integrity of cloned DNA was verified by sequencing analysis. All recombinant 6His-, GST- or MBP-fusion proteins were expressed in *Escherichia coli* BL21 (DE3) cells. The recombinant proteins were purified by Ni^+^ sepharose 6 fast flow (GE Healthcare), glutathione Sepharose 4B (GE Healthcare) or Amylose Resin High Flow (New England Biolabs) respectively according to the manufacturers’ instructions.

### *In vitro* pulldown assay

The recombinant MBP-Slp1^Cdc20^ and MBP, or GST-Pmk1(wild-type or mutants) and GST proteins were produced in bacteria and purified as written above. To detect the interaction between bacterially expressed Slp1 and Pmk1 or Sty1 expressed in yeast, about 1 μg of MBP-Slp1^Cdc20^ or MBP immobilized on amylose resin was incubated with cleared cell lysates prepared from yeast strains *nda3-KM311 pmk1-6His-HA* or *nda3-KM311 sty1-6His-HA*, which were HU synchronized and arrested at prometaphase by being grown at 18 °C for 6 hr. To detect the direct interaction between Slp1 and Pmk1, about 1 μg of GST-Pmk1 or GST immobilized on glutathione Sepharose 4B resin was incubated with cleared bacterial lysates expressing MBP-Slp1^Cdc20^. Samples were incubated for 1-3 hr at 4 °C. Resins were washed 3 times with lysis buffer (6 mM Na_2_HPO_4_, 4 mM NaH_2_PO_4_, 1% NP-40, 150 mM NaCl, 2 mM EDTA, 50 mM NaF, 0.1 mM Na_3_VO_4_, plus protease inhibitors), suspended in SDS sample buffer, and then subject to SDS-PAGE electrophoresis and Coomassie brilliant blue (CBB) staining.

### Non-radioactive *in vitro* kinase assay using fission yeast proteins

MBP-Slp1^Cdc20^, GST-Slp1^Cdc20^, GST-Slp1^Cdc20^(1-190aa, 2A), GST-Slp1^Cdc20^(1-190aa, 4A), GST-Slp1^Cdc20^(1-190aa, 9A), GST-Mad2, GST-Mad2(11A), GST-Mad2(13A), 6His-Mad3, MBP-Atf1, GST-Atf1, GST-Pmk1, GST-Sty1, GST-Pek1^DD^, and GST-Wis1^DD^ fusions were expressed and purified from *Escherichia coli* with Ni^+^sepharose 6 fast flow, glutathione Sepharose 4B or amylose resin as described above. Once purified and after extensive washing, beads bound with GST-Pmk1, GST-Sty1, GST-Pek1(DD), GST-Wis1(DD) and fused substrates MBP-Slp1^Cdc20^, GST-Slp1^Cdc20^, GST-Slp1^Cdc20^(1-190aa, 2A), GST-Slp1^Cdc20^(1-190aa, 4A), GST-Slp1^Cdc20^(1-190aa, 9A), GST-Mad2, GST-Mad2(11A), GST-Mad2(13A), 6His-Mad3, MBP-Atf1 or GST-Atf1 were washed 3 times with kinase buffer (for Cdk1: 10 mM Tris-HCl (pH 7.4), 10 mM MgCl_2_, 1 mM DTT; for Pmk1: 50 mM Tris-HCl (pH 7.5), 10 mM MgCl_2_, 1 mM EGTA; for Sty1: 20 mM Tris-HCl (pH 8.0), 10 mM MgCl_2_), then the reactions were incubated in kinase buffer with 20 μM ATPγS (Sigma-Aldrich) at 30°C for 45 min. The kinase reaction was stopped by adding 20 mM EDTA, and the reaction mixture was alkylated after incubation at room temperature with 2.5 mM *p*-nitrobenzyl mesylate (PNBM) (Abcam) for 1 h. The alkylation reaction was stopped by boiling in SDS-PAGE loading buffer. Phosphorylated proteins were detected with rabbit monoclonal anti-Thiophosphate ester antibody (Abcam, ab239919).

### Identification of Mad2 phosphorylation sites

To map the potential phosphorylation sites of Mad2 which were phosphorylated by Sty1, *in vitro* kinase reaction was set up same as in the non-radioactive *in vitro* kinase assay except 20 μM ATP instead of ATP-γ-S was used. The band corresponding to GST-Mad2 was excised from Coomassie brilliant blue (CBB)-stained gel after SDS-PAGE and subjected to digestion. Tryptic peptides were extracted from gels using 0.15% formic acid, 67% acetonitrile and dried for analysis with an AB SCIEX Triple TOF 5600 system.

### Antibodies for immunoblotting

The following antibodies were used for immunoblot analyses: peroxidase-anti-peroxidase (PAP) soluble complex (Sigma-Aldrich; P1291); rabbit polyclonal anti-Myc (GeneScript; A00172-40); mouse monoclonal anti-GFP (Beijing Ray Antibody Biotech; RM1008); rat monoclonal anti-HA (Roche, Cat. No. 11 867 423 001); rabbit polyclonal anti-Slp1 (Kim et al., 1998); mouse monoclonal anti-p44/42 (Cell Signaling Technology; #4696S); rabbit polyclonal anti-phospho-p44/42 (detecting activated Pmk1 or Erk1/2) (Cell Signaling Technology; #9101); rabbit polyclonal anti-PSTAIRE (detecting Cdc2) (Santa Cruz Biotechnology; sc-53); rabbit monoclonal anti-Thiophosphate ester antibody (Abcam; ab239919); goat anti-GST HRP-conjugated antibody (RRID:AB_771429; GE Healthcare). Secondary antibodies used were goat anti-mouse or goat anti-rabbit polyclonal IgG (H+L) HRP conjugates (Thermo Fisher Scientific; #31430 or #32460).

### Fluorescence microscopy

Cdc13-GFP proteins were observed in cells after fixation with cold methanol. For DAPI staining of nuclei, cells were fixed with cold methanol, washed in PBS and resuspended in PBS plus 1 μg/ml DAPI. Photomicrographs of cells were obtained using a Nikon 80i fluorescence microscope coupled to a cooled CCD camera (Hamamatsu, ORCA-ER) or a Perkin Elmer spinning-disk confocal microscope (UltraVIEW^®^ VoX) with a 100x NA 1.49 TIRF oil immersion objective (Nikon) coupled to a cooled CCD camera (9100-50 EMCCD; Hamamatsu Photonics) and spinning disk head (CSU-X1, Yokogawa). Image processing, analysis and spindle length measurement were carried out using Element software (Nikon), ImageJ software (National Institutes of Health) and Adobe Photoshop.

### Statistical analyses and reproducibility

No statistical methods were used to predetermine sample size. Sample size was as large as practicable for all experiments. The experiments were randomized and the investigators were blinded to group allocation during experiments but not blinded to outcome assessment and data analysis.

All experiments were independently repeated two to more than three times with similar results obtained. For quantitative analyses of each time course experiment, 300 cells were counted for each time point or sample. The same sample was not measured or counted repeatedly. No data were excluded from our studies. Data collection and statistical analyses were performed using Microsoft Office Excel or GraphPad Prism 6 softwares. Data are expressed as mean values with error bars (± standard deviation/s.d.) and were compared using unpaired *t*-tests or two-tailed Student’s *t*-tests unless indicated otherwise.

## Data availability statement

Strains and plasmids are available upon request. The authors affirm that all data necessary for confirming the conclusions of the article are present within the article, figures, and tables.

## ACKNOWLEDGMENTS

We thank Kathy Gould, Silke Hauf, Jonathan Millar, Takashi Toda, Yoshinori Watanabe, Elena Hidalgo and National BioResource Project (NBRP), Japan (http://yeast.nig.ac.jp/yeast/) for fission yeast strains or plasmids; Yaying Wu, Zheni Xu and Chang-chuan Xie for help with mass spectrometry analysis; Qing-feng Liu, Xin Chen and Li-xin Hong for help with confocal microscopy imaging. This work was supported by grants from the National Natural Science Foundation of China (No. 32170731, No. 31171298, No. 31671411) to Q.W. Jin.

## Author Contributions

L.S., S.B., and D.D. performed all immunoblotting and co-immunoprecipitation experiments in fission yeast; J.C. performed all time course analyses on SAC activation and silencing and serial dilution spot assays with help from Z.L. and D.D.; Z.L., Q.J. and Y.W. conceived the study; Y.W. and Q.J. designed the experiments and supervised the research; Y.W. and Q.J. wrote the paper with the inputs from Z.L.

## References

Akera, T., Y. Goto, M. Sato, M. Yamamoto, and Y. Watanabe. 2015. Mad1 promotes chromosome congression by anchoring a kinesin motor to the kinetochore. Nat Cell Biol. 17:1124–1133.

Alfieri, C., L. Chang, Z. Zhang, J. Yang, S. Maslen, M. Skehel, and D. Barford. 2016. Molecular basis of APC/C regulation by the spindle assembly checkpoint. Nature. 536:431–436.

Bardwell, A.J., L.J. Flatauer, K. Matsukuma, J. Thorner, and L. Bardwell. 2001. A conserved docking site in MEKs mediates high-affinity binding to MAP kinases and cooperates with a scaffold protein to enhance signal transmission. J Biol Chem. 276:10374–10386.

Bardwell, L., and J. Thorner. 1996. A conserved motif at the amino termini of MEKs might mediate high-affinity interaction with the cognate MAPKs. Trends in biochemical sciences. 21:373–374.

Bulavin, D.V., Y. Higashimoto, I.J. Popoff, W.A. Gaarde, V. Basrur, O. Potapova, E. Appella, and A.J. Fornace, Jr. 2001. Initiation of a G2/M checkpoint after ultraviolet radiation requires p38 kinase. Nature. 411:102–107.

Cansado, J., T. Soto, A. Franco, J. Vicente-Soler, and M. Madrid. 2021. The Fission Yeast Cell Integrity Pathway: A Functional Hub for Cell Survival upon Stress and Beyond. J Fungi (Basel). 8.

Chao, W.C., K. Kulkarni, Z. Zhang, E.H. Kong, and D. Barford. 2012. Structure of the mitotic checkpoint complex. Nature. 484:208–213.

Chen, R.H. 2004. Phosphorylation and activation of Bub1 on unattached chromosomes facilitate the spindle checkpoint. EMBO J. 23:3113–3121.

Chen, Y.H., G.Y. Wang, H.C. Hao, C.J. Chao, Y. Wang, and Q.W. Jin. 2017. Facile manipulation of protein localization in fission yeast through binding of GFP-binding protein to GFP. Journal of Cell Science. 130:1003–1015.

Chung, E., and R.H. Chen. 2003. Phosphorylation of Cdc20 is required for its inhibition by the spindle checkpoint. Nat Cell Biol. 5:748–753.

Fujimitsu, K., M. Grimaldi, and H. Yamano. 2016. Cyclin-dependent kinase 1-dependent activation of APC/C ubiquitin ligase. Science. 352:1121–1124.

Gonzalez, F.A., D.L. Raden, and R.J. Davis. 1991. Identification of substrate recognition determinants for human ERK1 and ERK2 protein kinases. J Biol Chem. 266:22159–22163.

Hartmuth, S., and J. Petersen. 2009. Fission yeast Tor1 functions as part of TORC1 to control mitotic entry through the stress MAPK pathway following nutrient stress. J Cell Sci. 122:1737–1746.

Heinrich, S., E.M. Geissen, J. Kamenz, S. Trautmann, C. Widmer, P. Drewe, M. Knop, N. Radde, J. Hasenauer, and S. Hauf. 2013. Determinants of robustness in spindle assembly checkpoint signalling. Nat Cell Biol. 15:1328–1339.

Hiraoka, Y., T. Toda, and M. Yanagida. 1984. The NDA3 gene of fission yeast encodes beta-tubulin: a cold-sensitive nda3 mutation reversibly blocks spindle formation and chromosome movement in mitosis. Cell. 39:349–358.

Izawa, D., and J. Pines. 2015. The mitotic checkpoint complex binds a second CDC20 to inhibit active APC/C. Nature. 517:631–634.

Jia, L., B. Li, and H. Yu. 2016. The Bub1-Plk1 kinase complex promotes spindle checkpoint signalling through Cdc20 phosphorylation. Nature communications. 7:10818.

Kapanidou, M., N.L. Curtis, and V.M. Bolanos-Garcia. 2017. Cdc20: At the Crossroads between Chromosome Segregation and Mitotic Exit. Trends in biochemical sciences. 42:193–205.

Kim, D.U., J. Hayles, D. Kim, V. Wood, H.O. Park, M. Won, H.S. Yoo, T. Duhig, M. Nam, G. Palmer, S. Han, L. Jeffery, S.T. Baek, H. Lee, Y.S. Shim, M. Lee, L. Kim, K.S. Heo, E.J. Noh, A.R. Lee, Y.J. Jang, K.S. Chung, S.J. Choi, J.Y. Park, Y. Park, H.M. Kim, S.K. Park, H.J. Park, E.J. Kang, H.B. Kim, H.S. Kang, H.M. Park, K. Kim, K. Song, K.B. Song, P. Nurse, and K.L. Hoe. 2010. Analysis of a genome-wide set of gene deletions in the fission yeast Schizosaccharomyces pombe. Nat Biotechnol. 28:617–623.

Kim, S.H., D.P. Lin, S. Matsumoto, A. Kitazono, and T. Matsumoto. 1998. Fission yeast Slp1: an effector of the Mad2-dependent spindle checkpoint. Science. 279:1045–1047.

Kraft, C., F. Herzog, C. Gieffers, K. Mechtler, A. Hagting, J. Pines, and J.M. Peters. 2003. Mitotic regulation of the human anaphase-promoting complex by phosphorylation. EMBO J. 22:6598–6609.

Labit, H., K. Fujimitsu, N.S. Bayin, T. Takaki, J. Gannon, and H. Yamano. 2012. Dephosphorylation of Cdc20 is required for its C-box-dependent activation of the APC/C. EMBO J. 31:3351–3362.

Lopez-Aviles, S., M. Grande, M. Gonzalez, A.L. Helgesen, V. Alemany, M. Sanchez-Piris, O. Bachs, J.B. Millar, and R. Aligue. 2005. Inactivation of the Cdc25 phosphatase by the stress-activated Srk1 kinase in fission yeast. Mol Cell. 17:49–59.

Lopez-Aviles, S., E. Lambea, A. Moldon, M. Grande, A. Fajardo, M.A. Rodriguez-Gabriel, E. Hidalgo, and R. Aligue. 2008. Activation of Srk1 by the mitogen-activated protein kinase Sty1/Spc1 precedes its dissociation from the kinase and signals its degradation. Mol Biol Cell. 19:1670–1679.

Manke, I.A., A. Nguyen, D. Lim, M.Q. Stewart, A.E. Elia, and M.B. Yaffe. 2005. MAPKAP kinase-2 is a cell cycle checkpoint kinase that regulates the G2/M transition and S phase progression in response to UV irradiation. Mol Cell. 17:37–48.

Mansfeld, J., P. Collin, M.O. Collins, J.S. Choudhary, and J. Pines. 2011. APC15 drives the turnover of MCC-CDC20 to make the spindle assembly checkpoint responsive to kinetochore attachment. Nat Cell Biol. 13:1234–1243.

May, K.M., F. Paldi, and K.G. Hardwick. 2017. Fission Yeast Apc15 Stabilizes MCC-Cdc20-APC/C Complexes, Ensuring Efficient Cdc20 Ubiquitination and Checkpoint Arrest. Curr Biol. 27:1221–1228.

Minshull, J., H. Sun, N.K. Tonks, and A.W. Murray. 1994. A MAP kinase-dependent spindle assembly checkpoint in Xenopus egg extracts. Cell. 79:475–486.

Murray, A.W. 2011. A brief history of error. Nat Cell Biol. 13:1178–1182.

Musacchio, A. 2015. The Molecular Biology of Spindle Assembly Checkpoint Signaling Dynamics. Curr Biol. 25:R1002–1018.

Pan, J., and R.H. Chen. 2004. Spindle checkpoint regulates Cdc20p stability in Saccharomyces cerevisiae. Genes Dev. 18:1439–1451.

Perez, P., and J. Cansado. 2010. Cell integrity signaling and response to stress in fission yeast. Curr Protein Pept Sci. 11:680–692.

Peters, J.M. 2006. The anaphase promoting complex/cyclosome: a machine designed to destroy. Nat Rev Mol Cell Biol. 7:644–656.

Petersen, J., and I.M. Hagan. 2005. Polo kinase links the stress pathway to cell cycle control and tip growth in fission yeast. Nature. 435:507–512.

Petersen, J., and P. Nurse. 2007. TOR signalling regulates mitotic commitment through the stress MAP kinase pathway and the Polo and Cdc2 kinases. Nat Cell Biol. 9:1263–1272.

Plotnikov, A., E. Zehorai, S. Procaccia, and R. Seger. 2011. The MAPK cascades: signaling components, nuclear roles and mechanisms of nuclear translocation. Biochim Biophys Acta. 1813:1619–1633.

Primorac, I., and A. Musacchio. 2013. Panta rhei: the APC/C at steady state. J Cell Biol. 201:177–189.

Qiao, R., F. Weissmann, M. Yamaguchi, N.G. Brown, R. VanderLinden, R. Imre, M.A. Jarvis, M.R. Brunner, I.F. Davidson, G. Litos, D. Haselbach, K. Mechtler, H. Stark, B.A. Schulman, and J.M. Peters. 2016. Mechanism of APC/CCDC20 activation by mitotic phosphorylation. Proc Natl Acad Sci U S A. 113:E2570–2578.

Ronkina, N., and M. Gaestel. 2022. MAPK-Activated Protein Kinases: Servant or Partner? Annu Rev Biochem.

Saitoh, S., K. Takahashi, and M. Yanagida. 1997. Mis6, a fission yeast inner centromere protein, acts during G1/S and forms specialized chromatin required for equal segregation. Cell. 90:131–143.

Sczaniecka, M., A. Feoktistova, K.M. May, J.S. Chen, J. Blyth, K.L. Gould, and K.G. Hardwick. 2008. The spindle checkpoint functions of Mad3 and Mad2 depend on a Mad3 KEN box-mediated interaction with Cdc20-anaphase-promoting complex (APC/C). J Biol Chem. 283:23039–23047.

Sewart, K., and S. Hauf. 2017. Different Functionality of Cdc20 Binding Sites within the Mitotic Checkpoint Complex. Curr Biol. 27:1213–1220.

Shiozaki, K., and P. Russell. 1995. Cell-cycle control linked to extracellular environment by MAP kinase pathway in fission yeast. Nature. 378:739–743.

Shiozaki, K., M. Shiozaki, and P. Russell. 1998. Heat stress activates fission yeast Spc1/StyI MAPK by a MEKK-independent mechanism. Mol Biol Cell. 9:1339–1349.

Smith, D.A., W.M. Toone, D. Chen, J. Bahler, N. Jones, B.A. Morgan, and J. Quinn. 2002. The Srk1 protein kinase is a target for the Sty1 stress-activated MAPK in fission yeast. J Biol Chem. 277:33411–33421.

Steen, J.A., H. Steen, A. Georgi, K. Parker, M. Springer, M. Kirchner, F. Hamprecht, and M.W. Kirschner. 2008. Different phosphorylation states of the anaphase promoting complex in response to antimitotic drugs: a quantitative proteomic analysis. Proc Natl Acad Sci U S A. 105:6069–6074.

Sudakin, V., G.K. Chan, and T.J. Yen. 2001. Checkpoint inhibition of the APC/C in HeLa cells is mediated by a complex of BUBR1, BUB3, CDC20, and MAD2. J Cell Biol. 154:925–936.

Sugiura, R., T. Toda, S. Dhut, H. Shuntoh, and T. Kuno. 1999. The MAPK kinase Pek1 acts as a phosphorylation-dependent molecular switch. Nature. 399:479–483.

Sullivan, M., and D.O. Morgan. 2007. Finishing mitosis, one step at a time. Nat Rev Mol Cell Biol. 8:894–903.

Takenaka, K., Y. Gotoh, and E. Nishida. 1997. MAP kinase is required for the spindle assembly checkpoint but is dispensable for the normal M phase entry and exit in Xenopus egg cell cycle extracts. J Cell Biol. 136:1091–1097.

Trautmann, S., S. Rajagopalan, and D. McCollum. 2004. The S. pombe Cdc14-like phosphatase Clp1p regulates chromosome biorientation and interacts with Aurora kinase. Dev Cell. 7:755–762.

Uzunova, K., B.T. Dye, H. Schutz, R. Ladurner, G. Petzold, Y. Toyoda, M.A. Jarvis, N.G. Brown, I. Poser, M. Novatchkova, K. Mechtler, A.A. Hyman, H. Stark, B.A. Schulman, and J.M. Peters. 2012. APC15 mediates CDC20 autoubiquitylation by APC/C(MCC) and disassembly of the mitotic checkpoint complex. Nat Struct Mol Biol. 19:1116–1123.

Vanoosthuyse, V., and K.G. Hardwick. 2009. A novel protein phosphatase 1-dependent spindle checkpoint silencing mechanism. Curr Biol. 19:1176–1181.

Wang, Y., W.Z. Li, A.E. Johnson, Z.Q. Luo, X.L. Sun, A. Feoktistova, W.H. McDonald, I. McLeod, J.R. Yates, 3rd, K.L. Gould, D. McCollum, and Q.W. Jin. 2012. Dnt1 acts as a mitotic inhibitor of the spindle checkpoint protein dma1 in fission yeast. Mol Biol Cell. 23:3348–3356.

Yamaguchi, M., R. VanderLinden, F. Weissmann, R. Qiao, P. Dube, N.G. Brown, D. Haselbach, W. Zhang, S.S. Sidhu, J.M. Peters, H. Stark, and B.A. Schulman. 2016. Cryo-EM of Mitotic Checkpoint Complex-Bound APC/C Reveals Reciprocal and Conformational Regulation of Ubiquitin Ligation. Mol Cell. 63:593–607.

Yoon, H.J., A. Feoktistova, B.A. Wolfe, J.L. Jennings, A.J. Link, and K.L. Gould. 2002. Proteomics analysis identifies new components of the fission and budding yeast anaphase-promoting complexes. Curr Biol. 12:2048–2054.

Yu, H. 2007. Cdc20: a WD40 activator for a cell cycle degradation machine. Mol Cell. 27:3–16.

Zhang, S., L. Chang, C. Alfieri, Z. Zhang, J. Yang, S. Maslen, M. Skehel, and D. Barford. 2016. Molecular mechanism of APC/C activation by mitotic phosphorylation. Nature. 533:260–264.

Zhao, Y., and R.H. Chen. 2006. Mps1 phosphorylation by MAP kinase is required for kinetochore localization of spindle-checkpoint proteins. Curr Biol. 16:1764–1769.

Zich, J., K. May, K. Paraskevopoulos, O. Sen, H.M. Syred, S. van der Sar, H. Patel, J.J. Moresco, A. Sarkeshik, J.R. Yates, 3rd, J. Rappsilber, and K.G. Hardwick. 2016. Mps1Mph1 Kinase Phosphorylates Mad3 to Inhibit Cdc20Slp1-APC/C and Maintain Spindle Checkpoint Arrests. PLoS Genet. 12:e1005834.

Zich, J., A.M. Sochaj, H.M. Syred, L. Milne, A.G. Cook, H. Ohkura, J. Rappsilber, and K.G. Hardwick. 2012. Kinase activity of fission yeast Mph1 is required for Mad2 and Mad3 to stably bind the anaphase promoting complex. Curr Biol. 22:296–301.

Zuin, A., M. Carmona, I. Morales-Ivorra, N. Gabrielli, A.P. Vivancos, J. Ayte, and E. Hidalgo. 2010. Lifespan extension by calorie restriction relies on the Sty1 MAP kinase stress pathway. EMBO J. 29:981–991.

