## Supplementary Information for "A dual mechanism of APC/C inhibition by MAP kinases"

**This PDF file includes:**

Figures S1 to S10

Table S1

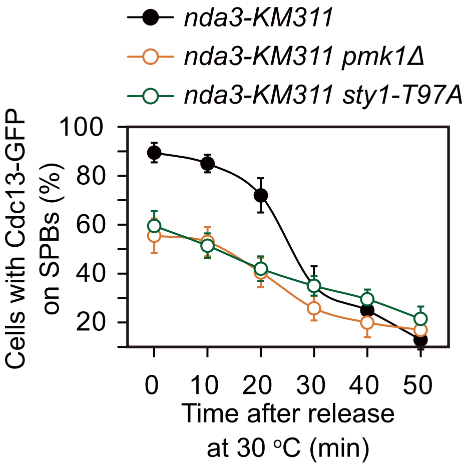
nda3-KM311
nda3-KM311 pmk1Δ
nda3-KM311 sty1-T97A
Cells with Cdc13-GFP on SPBs (%)
100
80
60
40
20
0
10
20
30
40
50
Time after release at 30 °C (min)

**Supplemental Figure legends:**

**Supplemental Figure S1. Related to Figure 1D.**

**Inactivation of CIP or SAP signaling by *pmk1*Δ or *sty1-T97A* compromises full SAC activation.**

Full time course analysis on SAC activation and silencing. Cells carrying Cdc13-GFP were grown at the permissive temperature for *nda3-KM311* (30 ℃) to mid-log phase, synchronized at S phase by HU. Cells were then washed and released at the restrictive temperature (18 ℃) for 6 hours and finally shifted back to the permissive temperature 30 ℃. Samples were collected at 10 min intervals and then subjected to microscopy analyses. *sty1-T97A* was inactivated by 5μM 3-BrB-PP1.

Sun et al., Supplemental Figure S2.

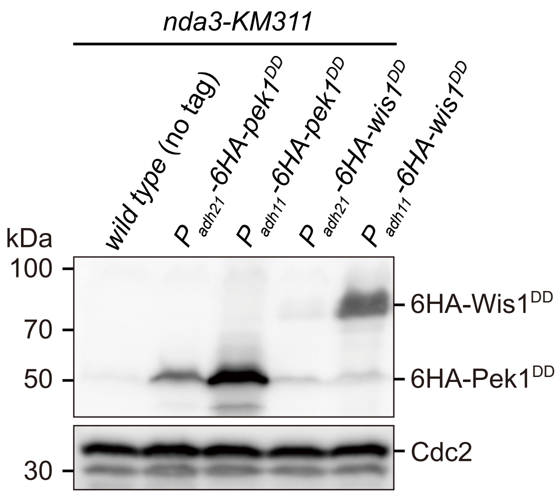

**Supplemental Figure S2. Examination of protein levels of overexpressed constitutively active MAPKKs (MAP kinase kinases) Pek1$^{DD}$ and Wis1$^{DD}$.**

Cells with indicated genotypes were grown at the permissive temperature for *nda3-KM311* (30 ℃) to mid-log phase and then arrested at 18 ℃ for 6 hours. Protein samples were prepared and immunoblotting was performed with anti-HA and anti-Cdc2 antibodies to detect total 6HA-Pek1$^{DD}$, 6HA-Wis1$^{DD}$ and Cdc2, respectively. Note $P_{adh11}$ is a stronger version of $P_{adh21}$ promoter.

Sun et al., Supplemental Figure S3.

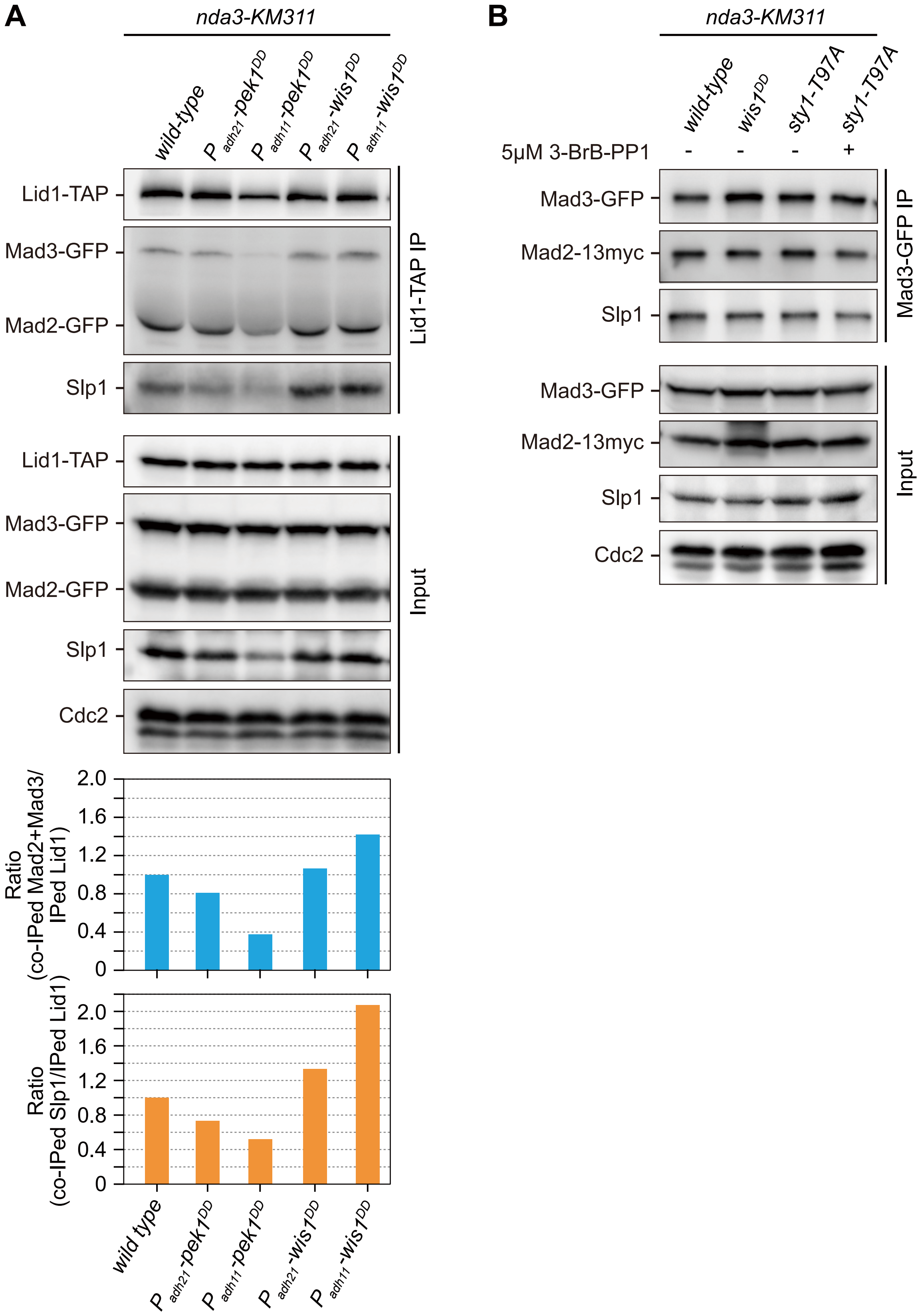

**Supplemental Figure S3. Related to Figure 2A.**

**Constitutive activation of Wis1-Sty1 but not Pek1-Pmk1 signaling enhances the association of MCC with APC/C, and SAP signaling does not affect MCC assembly.**

(**A**) Cells with indicated genotypes were grown at 30 ℃ to mid-log phase and arrested at 18 ℃ for 6 hours. The association of Mad2, Mad3 and Slp1$^{Cdc20}$ to Lid1 was assessed by immunoprecipitation of Lid1-TAP and immunoblotting with peroxidase-anti-peroxidase (PAP), anti-GFP, anti-Slp1 and anti-PSTAIRE antibodies to detect Lid1-TAP, Mad2-GFP, Mad3-GFP, Slp1 and Cdc2, respectively. For samples overexpressing *wis1*$^{DD}$ or *pek1*$^{DD}$, the amount of co-immunoprecipitated Mad2, Mad3 and Slp1 was quantified by being normalized to those of total immunoprecipitated Lid1 in each sample, with the relative ratio between Mad2-GFP plus Mad3-GFP or Slp1 and Lid1-TAP in wild-type sample set as 1.0.

Note that more Mad2, Mad3 and Slp1$^{Cdc20}$ was co-immunoprecipitated in $P_{adh21}$-*wis1*$^{DD}$ and $P_{adh11}$-*wis1*$^{DD}$ cells but not in $P_{adh21}$-*pek1*$^{DD}$ and $P_{adh11}$-*pek1*$^{DD}$ cells compared to wild-type cells, although the amount of Slp1$^{Cdc20}$ was comparable to that in wild-type cells. Also the abundance of Slp1$^{Cdc20}$ was significantly reduced in $P_{adh11}$-*pek1*$^{DD}$ cells (see input).

Results are representative of two independent experiments.

(**B**) Cells with indicated genotypes were grown and arrested as in (**A**). *sty1-T97A* was inactivated by 5μM 3-BrB-PP1. The association between Mad2, Mad3 and Slp1$^{Cdc20}$ was assessed by immunoprecipitation of Mad3-GFP using GFP-Trap beads and immunoblotting with anti-GFP, anti-myc, anti-Slp1 and anti-PSTAIRE antibodies to detect Mad3-GFP, Mad2-13myc, Slp1 and Cdc2, respectively.

Results are representative of two independent experiments.

Sun et al., Supplemental Figure S4.

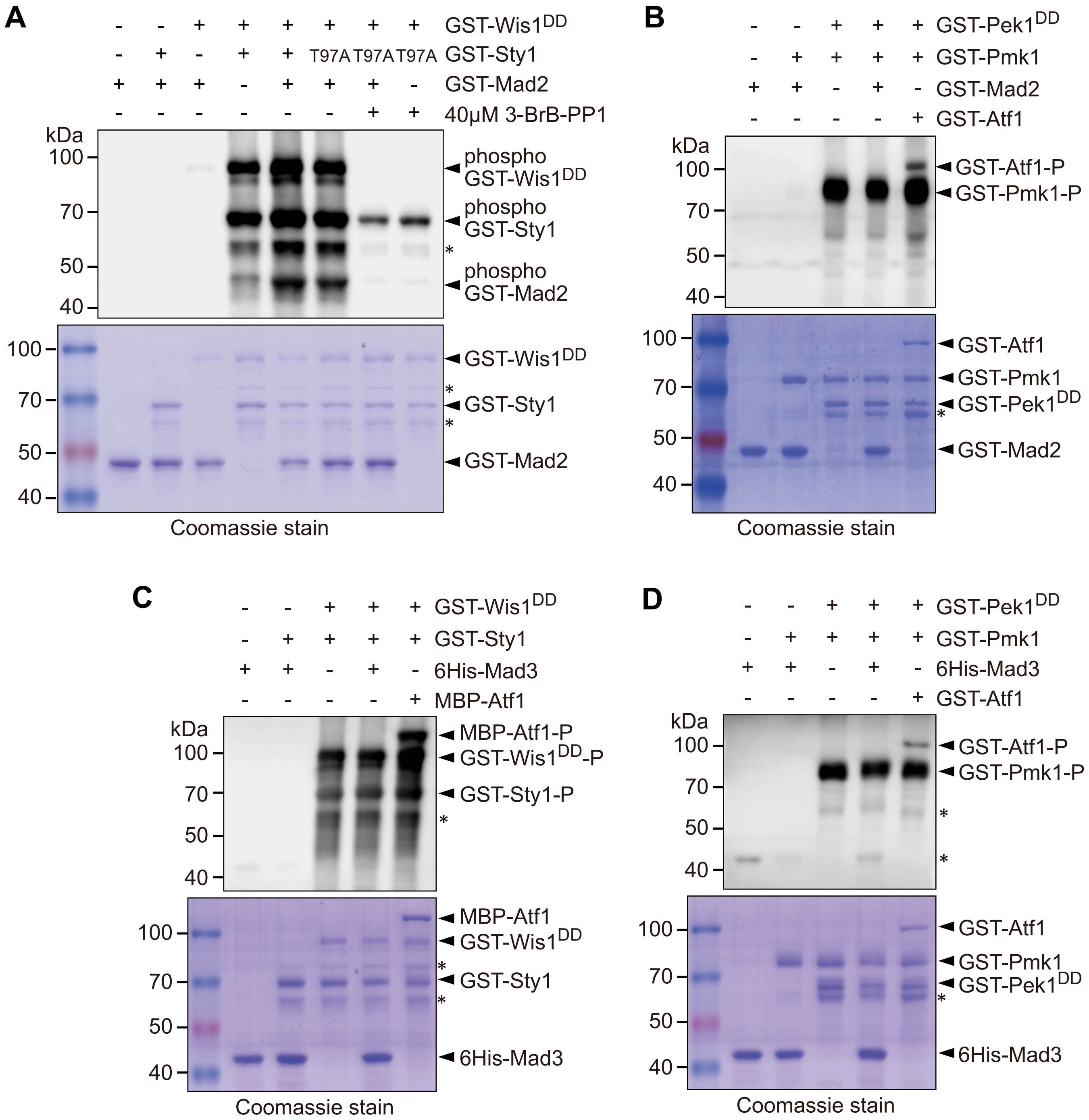

**Supplemental Figure S4. Recombinant Mad2 can be phosphorylated by Sty1 but not Pmk1 *in vitro*, and Mad3 cannot be phosphorylated by either MAPKs *in vitro*.**

*In vitro* non-radioactive kinase assay was performed with recombinant GST-Mad2 or 6His-Mad3 incubated with GST-Sty1 or GST-Sty1(T97A) (**A** and **C**), or GST-Pmk1 (**B** and **D**) in the presence or absence of GST-Wis1$^{DD}$ (**A** and **C**) or GST-Pek1$^{DD}$ (**B** and **D**). To inactivate Sty1(T97A) kinase activity, final concentration of 40μM ATP analog 3-BrB-PP1 was added into the reaction in (**A**). A known MAPK substrate Atf1 was used as a positive control. Note that a faint unspecific band was detected at roughly the same size as phosphorylated GST-Mad2 in (**A**).

Alignment of Mad2 homologues:

```
S.pombe          ---------MS-S-VPI-RTN-FSLKGSSKLVSEFFEYAVNSILFQRGIYPAEDFKVVRK 47
S.octosporus     ---------MA-A-VPI-RSN-LSLRGSSRLISEFFEYAVNSILFQRGIYPPEDFKVVRK 47
S.cryophilus     ---------MA-A-APV-RSN-LSLRGSSKVISEFFEYAVNSILFQRGIYPPEDFKVVRK 47
S.japonicus      --------MAT-T-VPT-RSS-LSLKGSAKLVSEFFEYAVNSILFQRGIYPPEDFKVVRK 48
S.cerevisiae     -----------MS-----QS--ISLKGSTRTVTEFFEYSINSILYQRGVYPAEDFVTVKK 42
T.ciferrii       --MSVAPPQES-SAAPT-RSN-VPLKGSAKLVAEFFEYSINSILYQRGIYPSEDFQVVRK 55
Y.lipolytica     -------MNKAPT-RDR-LSI----RGSSKTVAEFFEYSINTILYQRGIYPADDFQQVKK 47
P.jirovecii      ---------MTETNVAPTRSK-LSLKGSSKIVSEFFEYSINSILFQRGVYPADDFKIVKK 50
P.chlamydospora  ---------MAEAKADNEKTHKLSLKGSSKLVAEFFEYSINTILFQRGVYPAEDFSAVKK 51
H.sapiens        -------MALQLS-----REQGITLRGSAEIVAEFFSFGINSILYQRGIYPSETFTRVQK 48

S.pombe          YGLNMLVSVDEEVKTYIRKIVSQLHKWMFAKKIQKLILVITSKCSGEDLERWQFNVEMVD 107
S.octosporus     YGISMLVTVDEEVKTYIRKIISQLHRWTCGGKIQKVALVITNKDSGEDLERWQFNVEILQ 107
S.cryophilus     YGISMLVTVDEEVKTYIRKIILQLHRWTCGGKIQKVALVITNKDSGEDLERWQFNVEILQ 107
S.japonicus      YGINMLITIDDEVKAYIRRIIAQLHRWMYRGKIQKLVVVITDKDTGDDLERWQFNVEILC 108
S.cerevisiae     YDLTLLKTHDDELKDYIRKILLQVHRWLLGGKCNQLVLCIVDKDEGEVVERWSFNVQHI- 101
T.ciferrii       WDLNMLVTVDSNVKAYIKKIMSQLHKWLVGGKISKLVVVITSKDSGEVVERWQFDV---- 111
Y.lipolytica     YGINVLVTVDSEVKAYIRKIMGQLHKWLVGGKIQKLIVAITSKESGEVVERWQFDIHI-- 105
P.jirovecii      YGLNMLVTADTEVKAYIRKIMEQLHKWIEEGRISKLVIAIVSKDTLEVLERWQFDVQIFK 110
P.chlamydospora  YGLNMLVSSDDQVKAYIKKIMSQLNRWMVGGKISKLVVVITSKETGEHIERWQFDVQIF- 110
H.sapiens        YGLTLLVTTDLELIKYLNNVVEQLKDWLYKCSVQKLVVVISNIESGEVLERWQFDIECDK 108

S.pombe          ----TADQFQNIG------------NKEDELRVQKEIQALIRQITATVTFLPQL--EEQC 149
S.octosporus     --KDTPISPEN--------------TAQDDTRIQKEVQALIRQITATVTFLPQL--EGRC 149
S.cryophilus     --KDTTTVAE---------------HAQDESKIQKEVQALIRQITATVTFLPQL--EGRC 148
S.japonicus      KNEDSIGEE--------------SKEAKPEKEIQNEIQALIRQVTATITFLPQL--DTRC 152
S.cerevisiae     -SGNSNGQDD----------------VVDLNTTQSQIRALIRQITSSVTFLPELTKEGGY 144
T.ciferrii       ----SVMAGGGDGDS------QEGAAQKSQEDIQKEIQSIIRQITASVTFLPEL--QGRC 159
Y.lipolytica     SGKDTDTASKET--D-----TKTNGDKKSDDQIQQEIQAIMRQITASVSFLPVL--EDEC 156
P.jirovecii      NDNNENKENKLANEKNN------IATEKSEKEIHTEIQAIIRQITASVTFLPQL--EGQC 162
P.chlamydospora  SKSKSRSSSRKPATENAAPELDQIPTEKSEKEIQEEIQAIFRQITASVTFLPVL--DGNC 168
H.sapiens        ----TAKDDS-------------APREKSQKAIQDEIRSVIRQITATVTFLPLL--EVSC 149

S.pombe          TFNVLVYADKDSEVPTDWVDSDPRILRDAEQVQLRSFSTSMHKIDCQVAYRVNP-- 203
S.octosporus     TFNVLVYADRDSEVPTDWVDSDPRLIKNAEQVQLRSFSTNMHKIDCQVAYRFDP-- 203
S.cryophilus     TFNVLVYADRDSEVPTDWVDSDPRLIKNAEQVQLRSFSTSMHKIDCQVAYRFDP-- 202
S.japonicus      TFNVLVYADKDSEVPTDWVDSDPRQLQNAEQVQLRSFSTNMHKIDCQVAYRMN--- 205
S.cerevisiae     TFTVLAYTDADAKVPLEWADSNSKEIPDGEVVQFKTFSTNDHKVGAQVSYKY---- 196
T.ciferrii       TFNVLVYADGDADVPKEWGDSDPKDVKNAEQVQLRSFSTQSHKIDTLVAYKLGDD- 214
Y.lipolytica     TFNVLVYAEQDAPVPPEWADSANREIKNPEQVQLRSFSTNEHKIDTLVAYKLDR-- 210
P.jirovecii      TFNVLVYTDYSSEVPEEWGDSDAHEVDNAEQIKLRSFSTDLHKVDLQVAYKLADD- 217
P.chlamydospora  TFNVLVYADADSEVPVEWGDSDAKEIKNGEKVQLRSFSTTSHKVDTMVSYRLAD-- 222
H.sapiens        SFDLLIYTDKDLVVPEKWEESGPQFITNSEEVRLRSFTTTIHKVNSMVAYKIPVND 205
```

**Supplemental Figure S5. Conservation of potential MAPK phospho-sites within Mad2.**

Sequence alignment performed with *S. pombe* (*Schizosaccharomyces pombe*) Mad2 protein and its homologues in 9 other fungi species and humans: *Schizosaccharomyces octosporus*, *Schizosaccharomyces cryophilus*, *Schizosaccharomyces japonicas*, *Saccharomyces cerevisiae*, *Trichomonascus ciferrii*, *Yarrowia lipolytica*, *Pneumocystis jirovecii*, *Phaeomoniella chlamydospora*, and *Homo sapiens.* 13 potential Sty1 phosphorylation sites (Ser/Thr/Tyr) in *S. pombe* Mad2 identified by Mass spectrometry after *in vitro* kinase reaction and their conserved residues in other species are indicated in red and highlighted in yellow.

Sun et al., Supplemental Figure S6.

A

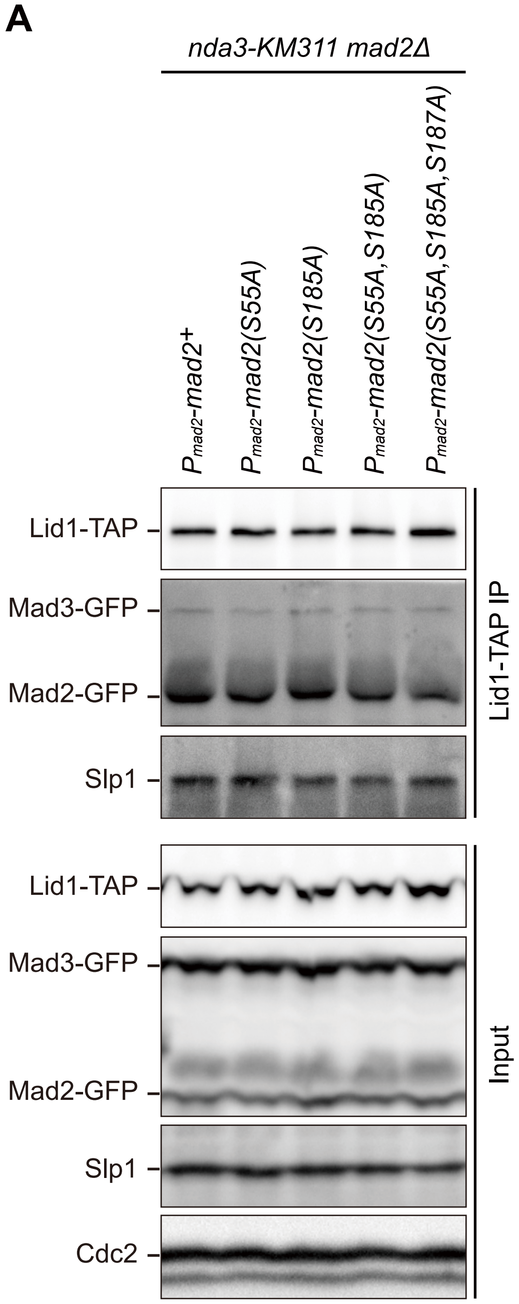


B

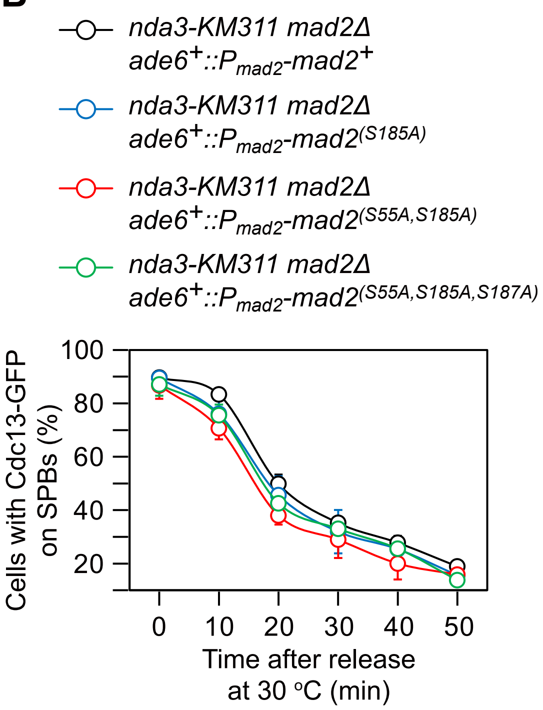

**Supplemental Figure S6. Individual or combined mutations of Ser55, Ser185 or Ser187 to alanine (A) are not sufficient to compromise MCC-APC/C association or SAC activation and maintenance.**

(**A**) Cells with indicated genotypes were grown at 30 ℃ to mid-log phase and arrested at 18 ℃ for 6 hours. The association of Mad2, Mad3 and Slp1$^{Cdc20}$ to Lid1 was assessed by immunoprecipitation of Lid1-TAP and immunoblotting with peroxidase-anti-peroxidase (PAP), anti-GFP, anti-Slp1 and anti-PSTAIRE antibodies to detect Lid1-TAP, Mad2-GFP, Mad3-GFP, Slp1 and Cdc2, respectively.

(**B**) Cells of indicated strains bearing Cdc13-GFP were grown at 30 ℃ to mid-log phase and arrested at 18 ℃ for 6 hours, and then released at 30 ℃. The percentage of cells with Cdc13-GFP on SPBs was assessed at each time point before or after release in each strain. Similar percentages of cells with Cdc13-GFP on SPBs in wild-type and *mad2* mutant strains indicate intact SAC activation when Ser55, Ser185 or Ser187 in Mad2 were mutated to alanine.

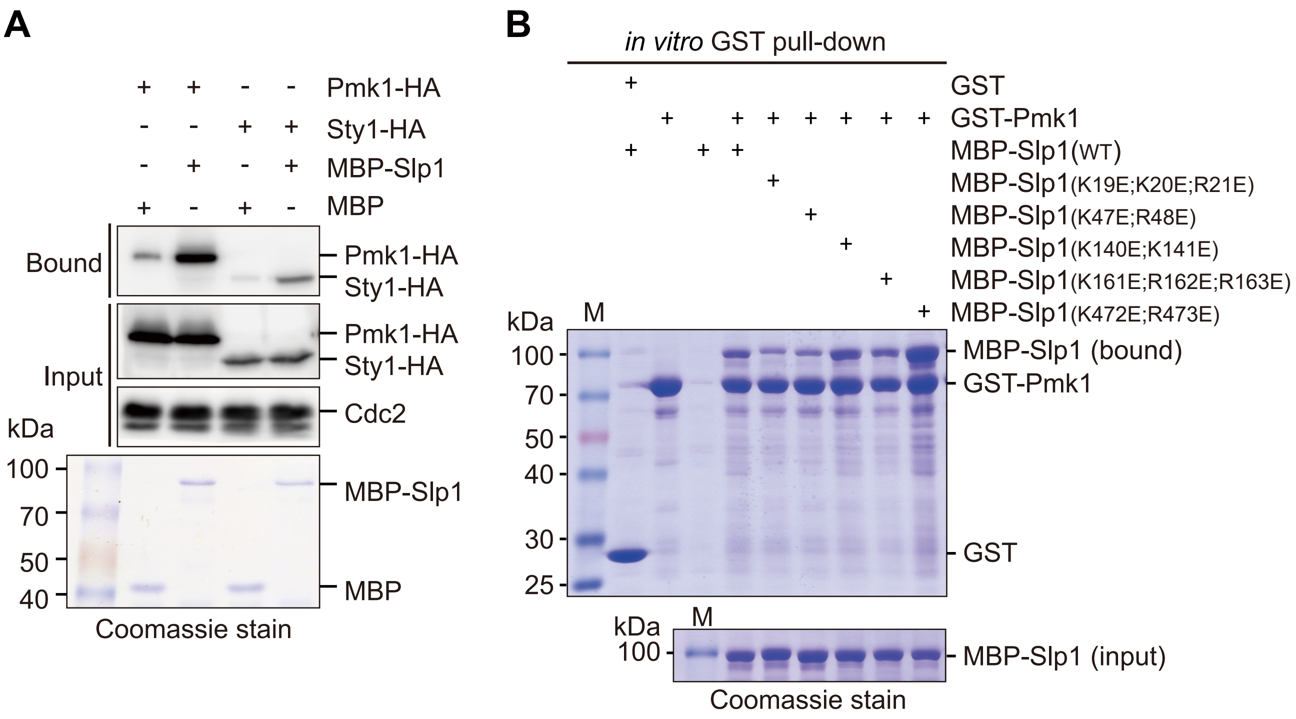
A
Pmk1-HA
Sty1-HA
MBP-Slp1
MBP
Bound
Input
Pmk1-HA
Sty1-HA
Pmk1-HA
Sty1-HA
Cdc2
kDa
100
70
50
40
MBP-Slp1
MBP
Coomassie stain
B
in vitro GST pull-down
GST
GST-Pmk1
MBP-Slp1(WT)
MBP-Slp1(K19E;K20E;R21E)
MBP-Slp1(K47E;R48E)
MBP-Slp1(K140E;K141E)
MBP-Slp1(K161E;R162E;R163E)
MBP-Slp1(K472E;R473E)
kDa
M
100
70
50
40
30
25
MBP-Slp1 (bound)
GST-Pmk1
GST
kDa
M
100
MBP-Slp1 (input)
Coomassie stain

**Supplemental Figure S7. Related to Figure 4B.**

**Pmk1 but not Sty1 directly binds to Slp1.**

(**A**) Two MAPKs Pmk1 and Sty1 bind $Slp1^{Cdc20}$ *in vitro*. *In vitro* binding assay was performed by incubating bacterially expressed MBP-$Slp1^{Cdc20}$ and yeast lysates prepared from *nda3-KM311 pmk1-HA-6His* or *nda3-KM311 sty1-HA-6His* cells, which were grown at 30 ℃ to mid-log phase and arrested at 18 ℃ for 6 hours. Proteins bound to beads with immobilized MBP or MBP-$Slp1^{Cdc20}$ were separated on SDS-PAGE and detected by Western blotting. Note that weak bands detected in MBP samples were due to unspecific background binding to beads. Coomassie blue staining shows inputs for MBP and MBP-$Slp1^{Cdc20}$.

(**B**) Examination on mutations of basic-residue patches within N-terminus of Slp1 disrupting the direct interaction between Slp1 and Pmk1. *In vitro* GST pull-down assays were performed with bacterially expressed GST-Pmk1 and MBP-Slp1 with wild-type Slp1 or mutants harboring Lys/Arg (K/R) to Glu (E) mutations. Proteins bound on GST beads were separated on SDS-PAGE and visualized by Coomassie blue staining. Note that Slp1 harboring clustered mutations K19E, K20E and R21E, or K47E and R48E within two most N-terminal basic-residue patches compromised interaction between Slp1 and Pmk1.

**A**

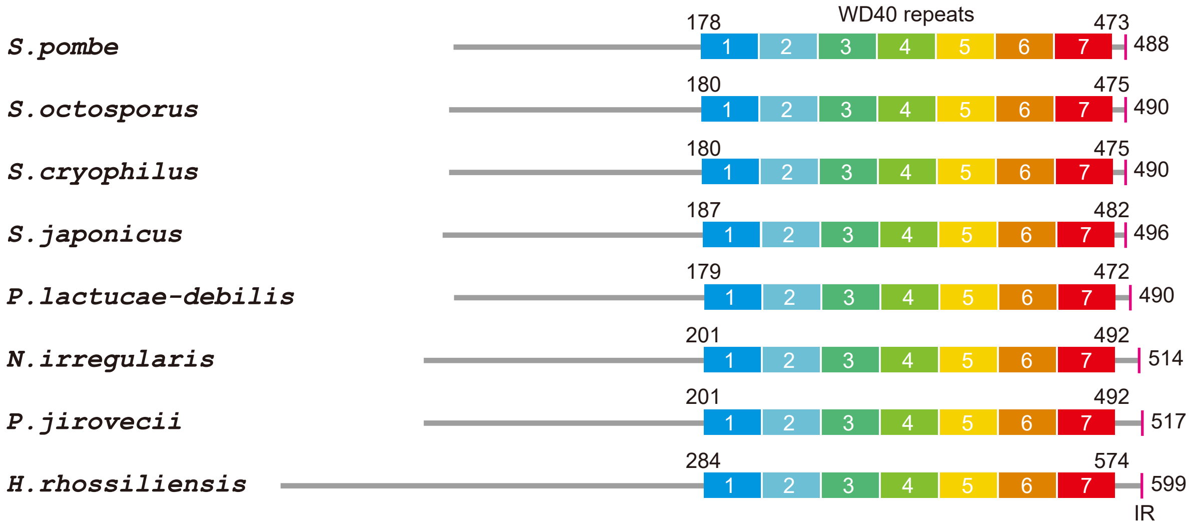


**B** Alignment of Slp1 homologues:

```
S.pombe             ----------MEIAGNSSTI---SPTFSTP--TKKRNLVFPN------SPITPLHQQA-- 37
S.octosporus        ----------MDVSSN-LSY---IPKFSTP--TKKRPLVYPN------SPITPLHQQA-- 36
S.cryophilus        ----------MDVSSN-LSY---VSKFSTP--TKKRPLVYPN------SPITPLHQQA-- 36
S.japonicus         ----------MDKKYLFSSS---AEECSTP--PRKRSYAFIS------SPATPLRQQV-- 37
P.lactucae-debilis  ------------------------MSLFTP---QRQSTSLSSILAGPGSPLTPRNVNI-- 31
N.irregularis       ----MFSTPVKQSAR--------CSLPRTPVSGKK--------LNFGASPLTPRRNAS-- 38
P.jirovecii         ----MLSTPMKHLDK--------YPVSCTPTSSIK----------GQKSPLTPRQTAA-- 36
H.rhossiliensis     MATASVSTPIKSHKGLFSSRTAGGRMPLTP--SPRQASTTTVPVNLNSSPFSPEGRATND 58

S.pombe             ----------------LLGRNGRSSKRC-SPKSSFIRN---SP-KIDVVNTDWSIPLC-- 74
S.octosporus        ----------------LLGKVSRSARK--SPTKVSSNFVRGSP-KLDVVSSDWSLSTSV- 76
S.cryophilus        ----------------LLGKVSRSAKK--SPTKTSFNFVRGSP-KLDVVASDWSLPTAV- 76
S.japonicus         ----------------LAGHVSRSALKN-SPKASVVIN---SP-KIEVVKRDWNVPTT-- 74
P.lactucae-debilis  ---------------------ARQLARQATPKSTLKRA---SPFKLDRVASDWTLTGTG- 66
N.irregularis       ----------IVG------VPKSV--KH-TPKSSIYRS---SPLKLEVVHSDWALTGTGN 76
P.jirovecii         ------------VLS-----HLHSAKRVYSPKSSIYRQ---TPLRLDIVASDWTLVGTGP 76
H.rhossiliensis     CRGSSKSTYGGNLMAHLSRPASRASHKE-SPKSNIARGVQ-TPRKALELGVSDFALTGTG 116

S.pombe             ----GSPR-------------NKSRPASRSDRFIPSRPNTANAFVNSI------------ 105
S.octosporus        ----SSPR-------------AKHHCPNHSDRFIPSRQNSANAFVSSQ------------ 107
S.cryophilus        ----NSPR-------------AKHRCLNHSDRFIPSRQNSANAFVSNQ------------ 107
S.japonicus         ----GSPK-------------PKRRPVIGSDRFIPVRPNIDNAHINNTNN---------- 107
P.lactucae-debilis  -----TPNKHAK---------RQVKGHGAGDRFIGRANPATAKLN-AE------------ 99
N.irregularis       AASIQTPGR--------QTRKRVARSIAQADRFIPTRNASSTAASKIDSRP--------- 119
P.jirovecii         LQ---SPQKNP----------KRPASRAQCDRFIPQRTHPASTATSKIAYPAV------- 116
H.rhossiliensis     K----TPASSKSRKAPLRQKSNKTTLNYSGDRFIPNRGASSAIANSGSSKLSLSDRQRNK 172

S.pombe             LAFKLDAPEAKKPVDLRTQHNRPQRPVV---TPAKRRFNTTPERVLDAPGIIDDYLNLL 186
S.octosporus        LAFKPDAPESKKPVDLRAQYNRPQKAVV---SQTKRKFATTPERVLDAPGIVDDYLNLL 188
S.cryophilus        LAFKPDAPESKKPVDLRAQYNRPQKPVV---SQTKRKFATTPERVLDAPGIVDDYLNLL 188
S.japonicus         LAFKPAPPESRKPVDLRAQYNRPAKPVA---SQV-RRIMTTPERVLDAPGIVDDYLNLL 195
P.lactucae-debilis  LAFKPEAPESKAPVLLNAQYNRPLRPVNT--ATVKRRINTVPERVLDAPGLIDDYLNLL 187
N.irregularis       LAFKPEAPESSRPVDLRSQYNRPLKPAAQ-AAQFRRRVLTAPERVLDAPGIVDDYLNLL 209
P.jirovecii         LAFKPTAPESSRPVDLRSQYNRPLKPAAL-NSQCRRRIATAPERVLDAPGLIDDYLNLL 209
H.rhossiliensis     LEFKPAPPESSKPIDLRQQYNRPLKPASTSSAQLRRRIATAPERVLDAPGLIDDYLNLL 292

S.pombe             LTKQVDIPAHDTRVLYSALSPDGRILSTAASDENLKFWRVYDGDHVKRPIPITKTPS--- 482
S.octosporus        VTKQVDIPAHDSRVLYSALSPDGRILSTAASDENLKFWRVNDSETKKRTGIFSKAGT--- 484
S.cryophilus        VTKQVDIPAHDSRVLYSALSPDGRILSTAASDENLKFWRVNDSETKKRTGIFAKAGP--- 484
S.japonicus         LVKQVDIPAHDTRVLYSSMSPDGCVLATAASDENLKFWKVYDNELKKKSV-VGKTSA--- 490
P.lactucae-debilis  LSKVVDIPAHDARVLHACLSPDGTTLATTSSDENLKFWKVFEA--KRRTAEH-----ERG 476
N.irregularis       LAKTVEISAHETRVLHSTLSPDGQVLATAAADENLKFWRVFEAA--KKSASGMADGKKS- 503
P.jirovecii         LVKSIDIPAHESRVLHSCLSPDGQVLATAASDENLKFWRVFEST--KKASCYVASISGSK 504
H.rhossiliensis     LVRNVEIPAHESRVLHSCLSPDGQMLATAAADESLKFWKVFE----KKAGAAGGIGGA-- 584
```

**Supplemental Figure S8. Conservation of putative MAPK phospho-sites and basic-residue patches within N-terminus of Slp1.**

(**A**) Schematic depiction of the *S. pombe* (*Schizosaccharomyces pombe*) Slp1 protein with its homologues in 7 other fungi species: *Schizosaccharomyces octosporus*, *Schizosaccharomyces cryophilus*, *Schizosaccharomyces japonicas*, *Protomyces lactucae-debilis*, *Neolecta irregularis*, *Pneumocystis jirovecii* and *Hirsutella rhossiliensis.* Positions of amino acids corresponding to seven WD40 repeats and IR motif are indicated.

(**B**) Local sequence alignment performed with *S. pombe* Slp1 and sequences from 7 other fungi species. 9 putative Pmk1 phosphorylation sites (SP/TP) in *S. pombe* Slp1 and their conserved positions in other species are indicated in red and highlighted in yellow. 5 potential basic-residue patches within N-terminus of Slp1 and their conserved positions in other species are highlighted in blue.

Sun et al., Supplemental Figure S9.

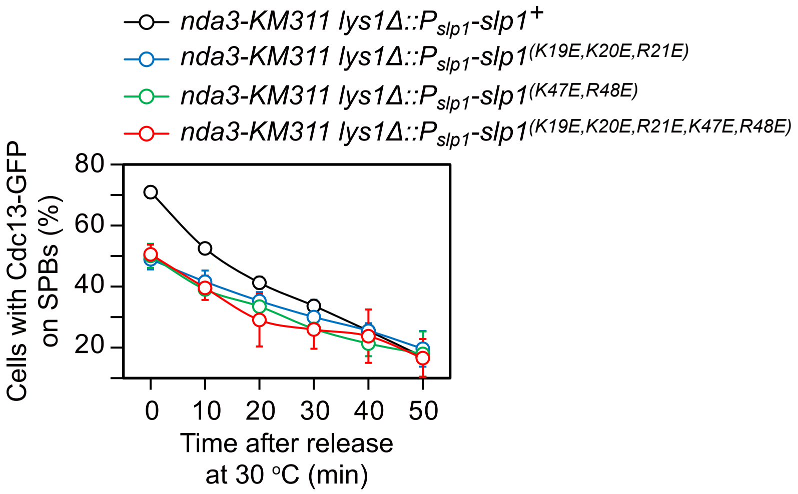

**Supplemental Figure S9. Related to Figure 4D.**

**Mutation of two basic-residue patches of Slp1 compromises SAC full activation and maintenance.**

Full time course analysis on SAC activation and silencing. Cells of indicated strains bearing Cdc13-GFP were grown at 30 ℃ to mid-log phase and arrested at 18 ℃ for 6 hours, and then the percentage of cells with Cdc13-GFP on SPBs was assessed right before and after release at 30 ℃ at each time point. Lowered percentages of cells with Cdc13-GFP on SPBs after being arrested at 18 ℃ for 6 hours (corresponding to 0 min time point in graph) in strains carrying mutations in two basic-residue patches of Slp1 indicate compromised SAC activation.

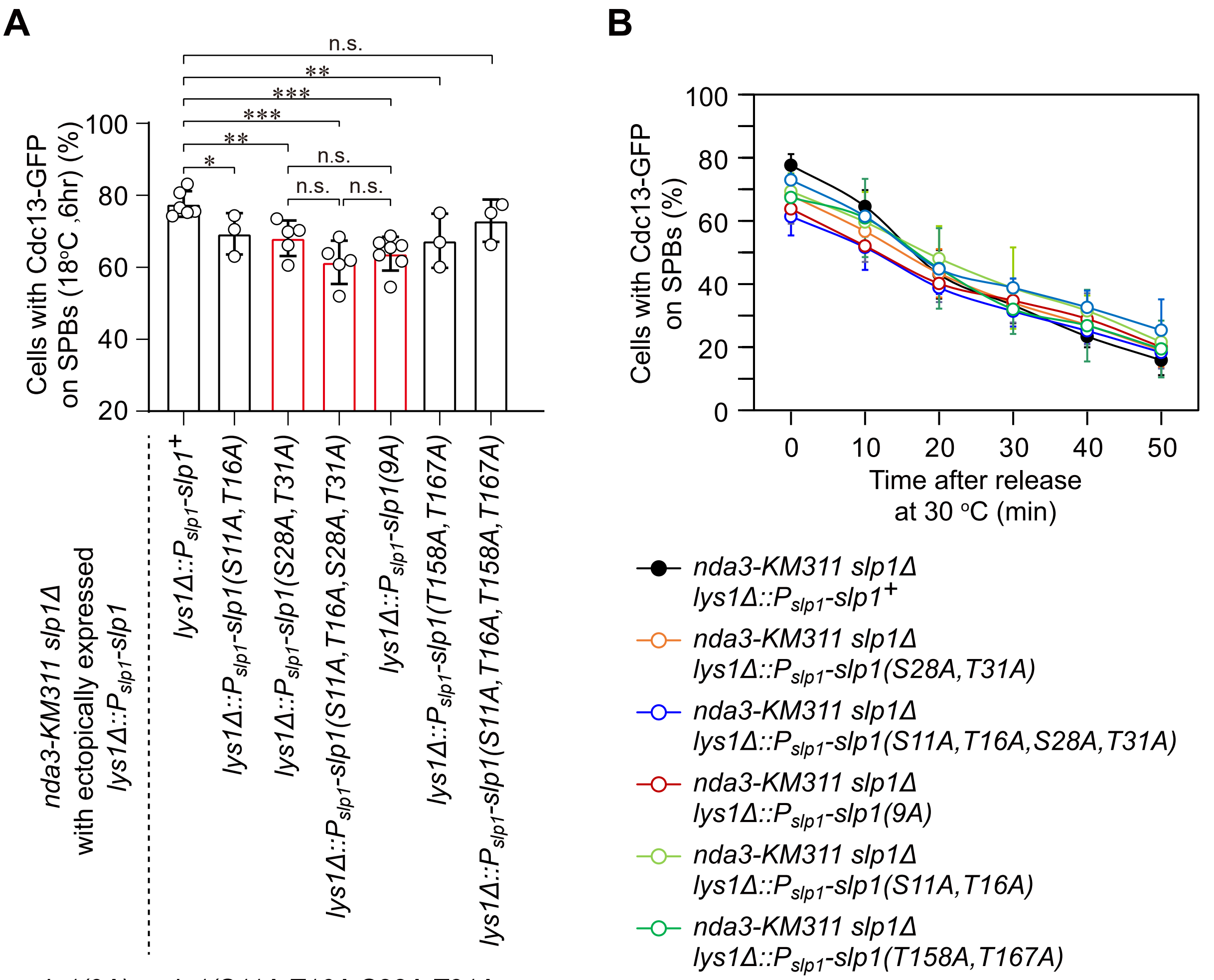


*slp1(9A) = slp1(S11A,T16A,S28A,T31A, S50A,S59A,S76A,T158A,T167A)*

**Supplemental Figure S10. Phosphorylation of Ser28 and Thr31 within N-terminus of Slp1 might play the major role in influencing APC/C activation.** Cells of indicated strains bearing *nda3-KM311* and Cdc13-GFP were grown at 30 ℃ to mid-log phase and arrested at 18 ℃ for 6 hours, and then the percentage of cells with Cdc13-GFP on SPBs was assessed right before (**A**) or after (**B**) release at 30 ℃, respectively. The data in (**A**) correspond to those at 0 min time point in graph (**B**).

**Supplemental Table S1. Yeast strains used in this study.**

| Strain | Genotype |
|---|---|
| JY3 | h$^{-}$ ade6-216 leu1-32 ura4-D18 |
| JY4 | h$^{+}$ ade6-216 leu1-32 ura4-D18 |
| JY227 | h$^{-}$ nda3-KM311 leu1-32 ura4-D18 ade6-21X |
| JY2162 | h$^{-}$ sty1Δ::ura4+ ura4-D18 leu1-32 |
| JY4773 | h$^{-}$ pmk1Δ ::ura4 ura4-D18 leu1-32 |
| JY4782 | h$^{-}$ spk1Δ::ura4 ura4-D18 leu1-32 |
| JY3058 | h$^{-}$ nda3-KM311 cdc13-117::cdc13-GFP:LEU2 ark1-as3::hyg$^{R}$ ura4- |
| JY4022 | h$^{?}$ nda3-KM311 lid1-TAP::kan$^{R}$ mad2-GFP::kan$^{R}$ mad3-GFP::his3$^{+}$ leu1-32 ura4-D18 his3-D1 ade6-216 |
| JY4023 | h$^{?}$ nda3-KM311 lid1-TAP::kan$^{R}$ mad2-GFP::kan$^{R}$ mad3-GFP::his3$^{+}$ leu1-32 ura4-D18 his3-D1 ade6-216 |
| JY8607 | h$^{+}$ nda3-KM311 cdc13-117::cdc13-GFP::LEU2 pDUAL-P$_{slp1(long2)}$-slp1-T$_{slp1}$::leu1$^{+}$ |
| JY8731 | h$^{-}$ nda3-KM311 cdc13-117::cdc13-GFP::LEU2 lys1Δ::pUC119-P$_{slp1(1504)}$-slp1-hyg$^{R}$ |
| JY8732 | h$^{-}$ nda3-KM311 cdc13-117::cdc13-GFP::LEU2 lys1Δ::pUC119-P$_{slp1(1504)}$-slp1-hyg$^{R}$ |
| JY77 | h$^{-}$ mad2Δ::ura4$^{+}$ u- l- |
| JY9182 | h$^{?}$ nda3-KM311 apc15Δ::kan$^{R}$ |
| JY9122 | h$^{?}$ nda3-KM311 apc15Δ::Kan$^{R}$ lid1-TAP-kan$^{R}$ mad2-GFP::kan$^{R}$ mad3-GFP-his3$^{+}$ |
| JY9371 | h$^{?}$ nda3-KM311 lid1-TAP-kan$^{R}$ mad2-GFP::kan$^{R}$ mad3-GFP-his3$^{+}$ pmk1Δ::ura4$^{+}$ |
| JY9241 | h$^{-}$ nda3-KM311 pmk1-HA-6His::ura4$^{+}$ ura4-D18 leu1-32 |
| JY9212 | h$^{+}$ nda3-KM311 sty1-3HA-kanMX |
| JY9245 | h$^{?}$ nda3-KM311 pmk1Δ::ura4$^{+}$ |
| JY9401 | h$^{-}$ nda3-KM311 sty1-T97A |
| JY9655 | h$^{?}$ nda3-KM311 pmk1Δ::ura4$^{+}$ sty1-T97A |
| JY9668 | h$^{?}$ nda3-KM311 lys1Δ::P$_{adh21}$-6×HA-pek1(DD)[(S234D, T238D)]::hyg$^{R}$ |
| JY9672 | h$^{?}$ nda3-KM311 lys1Δ::P$_{adh11}$-6×HA-pek1(DD)[(S234D, T238D)]::hyg$^{R}$ |
| JY9914 | h$^{-}$ nda3-KM311 lys1Δ::P$_{adh21}$-6×HA-wis1(DD)[S469D;T473D]::hyg$^{R}$ |
| JY9915 | h$^{-}$ nda3-KM311 lys1Δ::P$_{adh11}$-6×HA-wis1(DD)[S469D;T473D]::hyg$^{R}$ |
| JY9840 | h$^{+}$ nda3-KM311 lid1-TAP-kan$^{R}$ mad2-GFP::kan$^{R}$ mad3-GFP-his3$^{+}$ lys1Δ::P$_{adh21}$-6×HA-pek1(DD)[(S234D, T238D)]::hyg$^{R}$ |
| JY9860 | h$^{+}$ nda3-KM311 lid1-TAP-kan$^{R}$ mad2-GFP::kan$^{R}$ mad3-GFP-his3$^{+}$ lys1Δ::P$_{adh11}$-6×HA-pek1(DD)[(S234D, T238D)]::hyg$^{R}$ |
| JY9916 | h$^{-}$ nda3-KM311 lid1-TAP-kan$^{R}$ mad2-GFP::kan$^{R}$ mad3-GFP-his3$^{+}$ lys1Δ::P$_{adh21}$-6×HA-wis1(DD)[S469D;T473D]::hyg$^{R}$ |
| JY9934 | h- nda3-KM311 lid1-TAP-kan$^{R}$ mad2-GFP::kan$^{R}$ mad3-GFP-his3$^{+}$ lys1Δ::P$_{adh11}$-6×HA-wis1(DD)[S469D;T473D]::hyg$^{R}$ |
| JY9854 | h$^{?}$ nda3-KM311 lid1-TAP-kan$^{R}$ mad2-GFP::kan$^{R}$ mad3-GFP-his3$^{+}$ slp1Δ::ura4$^{+}$ lys1::pUC119-P$_{slp1(1504)}$-slp1(9A)::hyg$^{R}$ |
| JY9855 | h$^{?}$ nda3-KM311 lid1-TAP::kan$^{R}$ mad2-GFP- kan$^{R}$ mad3-GFP-his3$^{+}$ slp1Δ::ura4$^{+}$ lys1::pUC119-P$_{slp1(1504)}$-slp1(9E)::hyg$^{R}$ |
| JY5407 | h$^{+}$ nda3-KM311 cdc13-117::cdc13-GFP::LEU2 ura4$^{-}$ |
| JY9589 | h$^{+}$ nda3-KM311 cdc13-117::cdc13-GFP::LEU2 lys1Δ::P$_{adh11}$-6×HA-pek1(DD)[(S234D,T238D)]::hyg$^{R}$ |
| JY9590 | h$^{+}$ nda3-KM311 cdc13-117::cdc13-GFP::LEU2 |

| | |
|---|---|
| | lys1Δ::P$_{adh21}$-6×HA-pek1(DD)[(S234D,T238D)]::hyg$^{R}$ |
| JY9928 | h$^{+}$ nda3-KM311 cdc13-117::cdc13-GFP::LEU2<br>lys1Δ::P$_{adh21}$-6×HA-wis1(DD)[(S469D,T473D)]::hyg$^{R}$ ura4$^{-}$ |
| JY9929 | h$^{+}$ nda3-KM311 cdc13-117::cdc13-GFP::LEU2<br>lys1Δ::P$_{adh11}$-6×HA-wis1(DD)[(S469D,T473D)]::hyg$^{R}$ ura4$^{-}$ |
| JY9586 | h$^{+}$ lys1Δ::P$_{adh21}$-6×HA-pek1(DD)[(S234D, T238D)]::hyg$^{R}$ ura4-D18 leu1-32 |
| JY9585 | h$^{+}$ lys1Δ::P$_{adh11}$-6×HA-pek1(DD)[(S234D, T238D)]::hyg$^{R}$ ura4-D18 leu1-32 |
| JY9955 | h$^{+}$ lys1Δ::P$_{adh21}$-6×HA-wis1-DD[(S469D, T473D)]::hyg$^{R}$ ura4-D18 leu1-32 |
| JY9957 | h$^{+}$ lys1Δ::P$_{adh11}$-6×HA-wis1-DD[(S469D, T473D)]::hyg$^{R}$ ura4-D18 leu1-32 |
| JY9790 | h$^{-}$ nda3-KM311 lid1-TAP-kan$^{R}$ mad2-GFP::kan$^{R}$ mad3-GFP-his3$^{+}$ sty1-T97A |
| JY10123 | h$^{+}$ nda3-KM311 lid1-TAP-kan$^{R}$ mad2Δ::ura4$^{+}$ mad3-GFP-his3$^{+}$ |
| JY10251 | h$^{+}$ nda3-KM311 lid1-TAP-kan$^{R}$ mad2Δ::ura4$^{+}$ mad3-GFP-his3$^{+}$<br>ade6$^{+}$::P$_{adh21}$-mad2-GFP::nat$^{R}$ |
| JY10261 | h$^{-}$ nda3-KM311 lid1-TAP-kan$^{R}$ mad2Δ::ura4$^{+}$ mad3-GFP-his3$^{+}$<br>ade6$^{+}$::P$_{adh21}$-mad2(13A)-GFP::nat$^{R}$ |
| JY10271 | h$^{?}$ nda3-KM311 lid1-TAP-kan$^{R}$ mad3-GFP-his3$^{+}$ mad2Δ::ura4$^{+}$<br>ade6$^{+}$::P$_{adh21}$-mad2-GFP::nat$^{R}$ lys1Δ::P$_{adh11}$-6×HA-wis1(DD)[S469D,T473D]::hyg$^{R}$ |
| JY10273 | h$^{?}$ nda3-KM311 lid1-TAP-kan$^{R}$ mad3-GFP-his3$^{+}$ mad2Δ::ura4$^{+}$<br>ade6$^{+}$::P$_{adh21}$-mad2(13A)-GFP::nat$^{R}$<br>lys1Δ::P$_{adh11}$-6×HA-wis1(DD)[(S469D,T473D)]::hyg$^{R}$ |
| JY10193 | h$^{?}$ nda3-KM311 lid1-TAP-kan$^{R}$ mad3-GFP-his3$^{+}$ mad2Δ::ura4$^{+}$<br>Z::P$_{mad2(1238bp)}$-mad2-GFP::kan$^{R}$ his$^{+}$ |
| JY10264 | h$^{+}$ nda3-KM311 lid1-TAP-kan$^{R}$ mad2Δ::ura4$^{+}$ mad3-GFP-his3$^{+}$<br>ade6$^{+}$::P$_{mad2(1238bp)}$-mad2(S55A)-GFP::nat$^{R}$ |
| JY10252 | h$^{+}$ nda3-KM311 lid1-TAP-kan$^{R}$ mad2Δ::ura4$^{+}$ mad3-GFP-his3$^{+}$<br>ade6$^{+}$::P$_{mad2(1238bp)}$-mad2(S185A)-GFP::nat$^{R}$ |
| JY10253 | h$^{+}$ nda3-KM311 lid1-TAP-kan$^{R}$ mad2Δ::ura4$^{+}$ mad3-GFP-his3$^{+}$<br>ade6$^{+}$::P$_{mad2(1238bp)}$-mad2(S55A,S185A)-GFP::nat$^{R}$ |
| JY10254 | h$^{+}$ nda3-KM311 lid1-TAP-kan$^{R}$ mad2Δ::ura4$^{+}$ mad3-GFP-his3$^{+}$<br>ade6$^{+}$::P$_{mad2(1238bp)}$-mad2(S55A,S185A,S187A)-GFP::nat$^{R}$ |
| JY10104 | h$^{?}$ nda3-KM311 slp1Δ::ura4$^{+}$ lys1Δ::pUC119-P$_{slp1(1504)}$-slp1::hyg$^{R}$<br>cdc13-117::cdc13-GFP::LEU2<br>Z::P$_{adh11}$-6×HA-pek1(DD)[(S234D,T238D)]::hyg$^{R}$ |
| JY10103 | h$^{?}$ nda3-KM311 slp1Δ::ura4$^{+}$ lys1Δ::pUC119<br>P$_{slp1(1504)}$-slp1(S28A,T31A,S11A,T16A,S50A,S59A,S76A, T158A,T167A) [i.e.<br>slp1(9A)]::hyg$^{R}$ cdc13-117::cdc13-GFP::LEU2<br>Z::P$_{adh11}$-6×HA-pek1(DD)[(S234D,T238D)]::hyg$^{R}$ |
| JY10135 | h$^{?}$ nda3-KM311 slp1Δ::ura4$^{+}$ lys1Δ::pUC119-Pslp1(1504)-slp1::hygR<br>cdc13-117::cdc13-GFP::LEU2<br>ura4:: P$_{adh11}$-6×HA-wis1(DD)[(S469D,T473D)]::hyg$^{R}$ |
| JY10114 | h$^{?}$ nda3-KM311 slp1Δ::ura4$^{+}$ lys1Δ::pUC119-<br>P$_{slp1(1504)}$-slp1(S28A,T31A,S11A,T16A,S50A,S59A,S76A, T158A,T167A) [i.e.<br>slp1(9A)]::hyg$^{R}$ cdc13-117::cdc13-GFP::LEU2<br>ura4:: P$_{adh11}$-6×HA-wis1(DD)[(S469D,T473D)]::hyg$^{R}$ |
| JY10094 | h$^{?}$ nda3-KM311 apc15Δ::kan$^{R}$ |

| | |
|---|---|
| | cdc13-117::cdc13-GFP::LEU2 |
| JY10080 | $h^{?}$ nda3-KM311 apc15Δ::$kan^{R}$ cdc13-117::cdc13-GFP::LEU2 lys1Δ:: $P_{adh11}$-6×HA-pek1(DD)[(S234D,T238D)]::$hyg^{R}$ |
| JY10095 | $h^{?}$ nda3-KM311 apc15Δ::$kan^{R}$ cdc13-117::cdc13-GFP::LEU2 lys1Δ::$P_{adh11}$-6×HA-wis1(DD)[(S469D,T473D)]::$hyg^{R}$ |
| JY10106 | $h^{+}$ nda3-KM311 mad2Δ::$ura4^{+}$ cdc13-117::cdc13-GFP::LEU2 ura4- |
| JY10133 | $h^{-}$ nda3-KM311 mad2Δ::$ura4^{+}$ cdc13-117::cdc13-GFP::LEU2 lys1Δ::$P_{adh11}$-6×HA-wis1(DD)[(S469D,T473D)]::$hyg^{R}$ ura4- |
| JY10181 | $h^{+}$ nda3-KM311 mad2Δ::$ura4^{+}$ cdc13-117::cdc13-GFP::LEU2 Z::$P_{mad2}$-mad2::$kan^{R}$ ura4- |
| JY10175 | $h^{-}$ nda3-KM311 mad2Δ::$ura4^{+}$ cdc13-117::cdc13-GFP::LEU2 lys1Δ::$P_{adh11}$-6×HA-wis1(DD) [(S469D,T473D)]::$hyg^{R}$ Z::$P_{mad2}$-mad2::$kan^{R}$ ura4- |
| JY10180 | $h^{+}$ nda3-KM311 mad2Δ::$ura4^{+}$ cdc13-117::cdc13-GFP::LEU2 Z::$P_{mad2}$-mad2 (13A)::$kan^{R}$ ura4- |
| JY10214 | $h^{-}$ nda3-KM311 mad2Δ::$ura4^{+}$ cdc13-117::cdc13-GFP::LEU2 lys1Δ::$P_{adh11}$-6×HA-wis1(DD) [(S469D,T473D)]::$hyg^{R}$ Z::$P_{mad2}$-mad2(13A)::$kan^{R}$ ura4- |
| JY10259 | $h^{+}$ nda3-KM311 mad2Δ::$ura4^{+}$ cdc13-117::cdc13-GFP::LEU2 $ade6^{+}$::$P_{adh21}$-mad2-$nat^{R}$ |
| JY10279 | $h^{?}$ nda3-KM311 mad2Δ::$ura4^{+}$ cdc13-117::cdc13-GFP::LEU2 $ade6^{+}$::$P_{adh21}$-mad2-$nat^{R}$ lys1Δ::$P_{adh11}$-6×HA-wis1(DD)[S469D;T473D]::$hyg^{R}$ |
| JY10260 | $h^{+}$ nda3-KM311 mad2Δ::$ura4^{+}$ cdc13-117::cdc13-GFP::LEU2 $ade6^{+}$::$P_{adh21}$-mad2(13A)-$nat^{R}$ |
| JY10285 | $h^{?}$ nda3-KM311 mad2Δ::$ura4^{+}$ cdc13-117::cdc13-GFP::LEU2 $ade6^{+}$::$P_{adh21}$-mad2(13A)-$nat^{R}$ dnt1Δ::$kan^{R}$ |
| JY10281 | $h^{?}$ nda3-KM311 mad2Δ::$ura4^{+}$ cdc13-117::cdc13-GFP::LEU2 $ade6^{+}$::$P_{adh21}$-mad2(13A)-$nat^{R}$ lys1Δ::$P_{adh11}$-6×HA-wis1(DD)[S469D;T473D]::$hyg^{R}$ |
| JY9890 | $h^{-}$ nda3-KM311 slp1Δ::$ura4^{+}$ cdc13-117::cdc13-GFP::LEU2 lys1Δ::pUC119-$P_{slp1(1504)}$-slp1::$hyg^{R}$ ura4- |
| JY10366 | $h^{+}$ nda3-KM311 slp1Δ::$ura4^{+}$ cdc13-117::cdc13-GFP::LEU2 lys1Δ::pUC119-$P_{slp1(1504)}$-slp1(S28A,T31A)::$hyg^{R}$ |
| JY10364 | $h^{+}$ nda3-KM311 slp1Δ::$ura4^{+}$ cdc13-117::cdc13-GFP::LEU2 lys1Δ::pUC119-$P_{slp1(1504)}$-slp1(S28A,T31A,S11A,T16A)-$hyg^{R}$ ura4- leu1-32 |
| JY10358 | $h^{+}$ nda3-KM311 slp1Δ::$ura4^{+}$ cdc13-117::cdc13-GFP::LEU2 lys1Δ::pUC119-$P_{slp1(1504)}$-slp1(S11A,T16A,T167A, T158A)::$hyg^{R}$ |
| JY10365 | $h^{+}$ nda3-KM311 slp1Δ::$ura4^{+}$ cdc13-117::cdc13-GFP::LEU2 lys1Δ::pUC119-$P_{slp1(1504)}$-slp1(S11A,T16A)::$hyg^{R}$ |
| JY10367 | $h^{+}$ nda3-KM311 slp1Δ::$ura4^{+}$ cdc13-117::cdc13-GFP::LEU2 lys1Δ::pUC119-$P_{slp1(1504)}$-slp1(T158A, T167A)::$hyg^{R}$ |
| JY9892 | $h^{+}$ nda3-KM311 cdc13-117::cdc13-GFP::LEU2 lys1Δ::pUC119-$P_{slp1(1504)}$-slp1::$hyg^{R}$ ura4- |
| JY10452 | $h^{-}$ nda3-KM311 cdc13-117::cdc13-GFP::LEU2 lys1Δ::pUC119-$P_{slp1(1504)}$-slp1(K19E,K20E,R21E)-$hyg^{R}$ ura4- |

| | |
|---|---|
| JY10454 | *h*$^{+}$ *nda3-KM311 cdc13-117::cdc13-GFP::LEU2*<br>*lys1Δ::P*$_{slp1(1504)}$*-slp1(K47E,R48E)-hyg*$^{R}$ *ura4-* |
| JY10456 | *h*$^{-}$ *nda3-KM311 cdc13-117::cdc13-GFP::LEU2*<br>*lys1Δ::P*$_{slp1(1504)}$*-slp1(K19E,K20E,R21E,K47E,R48E)-hyg*$^{R}$ *ura4-* |
| JY10305 | *h*$^{+}$ *nda3-KM311 mad2Δ::ura4*$^{+}$ *cdc13-117::cdc13-GFP::LEU2*<br>*ade6*$^{+}$*::P*$_{mad2(1238bp)}$*-mad2-nat*$^{R}$ *ura4-* |
| JY10265 | *h*$^{+}$ *nda3-KM311 mad2Δ::ura4*$^{+}$ *cdc13-117::cdc13-GFP::LEU2*<br>*ade6*$^{+}$*::P*$_{mad2(1238bp)}$*-mad2(S185A)::nat*$^{R}$ *ura4-* |
| JY10266 | *h*$^{+}$ *nda3-KM311 mad2Δ::ura4*$^{+}$ *cdc13-117::cdc13-GFP::LEU2*<br>*ade6*$^{+}$*::P*$_{mad2(1238bp)}$*-mad2(S55A,S185A)::nat*$^{R}$ *ura4-* |
| JY10267 | *h*$^{+}$ *nda3-KM311 mad2Δ::ura4*$^{+}$ *cdc13-117::cdc13-GFP::LEU2*<br>*ade6*$^{+}$*::P*$_{mad2(1238bp)}$*-mad2(S55A,S185A,S187A)::nat*$^{R}$ *ura4-* |
| JY9777 | *h*$^{+}$ *nda3-KM311 slp1Δ::ura4*$^{+}$ *lys1Δ::pUC119-P*$_{slp1(1504)}$*-slp1-hyg*$^{R}$ *ura4-D18 leu1-32* |
| JY9890 | *h*$^{-}$ *nda3-KM311 slp1Δ::ura4*$^{+}$ *cdc13-117::cdc13-GFP::LEU2*<br>*lys1Δ::pUC119-P*$_{slp1(1504)}$*-slp1::hyg*$^{R}$ *ura4-* |
| JY9898 | *h*$^{+}$ *nda3-KM311 slp1Δ::ura4*$^{+}$ *pmk1Δ::ura4*$^{+}$ *sty1-T97A*<br>*lys1Δ::pUC119-P*$_{slp1(1504)}$*-slp1-hyg*$^{R}$ *cdc13-117::cdc13-GFP::LEU2 leu1- ura4-* |
| JY9177 | *h*$^{+}$ *nda3-KM311 mad3-GFP-his3*$^{+}$ *mad2-13myc::hygR* |
| JY9685 | *h*$^{+}$ *nda3-KM311 mad3-GFP-his3*$^{+}$ *mad2-13myc::hygR wis1-DD::ura4*$^{+}$ |
| JY9720 | *h*$^{+}$ *nda3-KM311 mad3-GFP-his3*$^{+}$ *mad2-13myc::hygR sty1-T97A* |
| JY9686 | *h*$^{-}$ *nda3-KM311 lid1-TAP-kan*$^{R}$ *mad2-GFP::kan*$^{R}$ *mad3-GFP-his3*$^{+}$ *apc15Δ::kan*$^{R}$<br>*lys1Δ::P*$_{adh11}$*-6×HA-wis1(DD)[S469D;T473D]::hyg*$^{R}$ |
